# ISG15-USP18 signaling restrains viperin-dependent metabolic antiviral restriction

**DOI:** 10.64898/2026.03.13.711570

**Authors:** Niklas L Kahler, Sefanit Rezene, Florine EM Scholte, Anoop T Ambikan, Elisa Saccon, Vanessa M Monteil, Prajakta Naval, Robin Macmillan, Simon Frick, Jenny Norén, Magda Lourda, Ali Mirazimi, Yenan T Bryceson, Ronaldo Lira-Junior, Vibhu Prasad, Ákos Végvári, Ujjwal Neogi, Connor GG Bamford, Anders Hofer, Eric Bergeron, Janne Purhonen, Soham Gupta

## Abstract

Type I interferon (IFN-I) responses are tightly regulated to balance antiviral defense with cellular homeostasis. In humans, interferon-stimulated gene 15 (ISG15) functions as a critical negative regulator of IFN-I signaling by stabilizing the IFN negative regulator USP18, yet the functional consequences of ISG15 deficiency remain elusive. Here, we show that the loss of ISG15 exaggerates the JAK-STAT activation and, downstream, amplifies multiple ISGs including the nucleotide-modifying enzyme RSAD2 (viperin). Our quantitative proteomics, genetic reconstitution, and signaling analyses establish that defective USP18 stabilization skews the IFN response towards viperin expression. This amplified ISG network promotes viperin-catalyzed accumulation of the antiviral nucleotide analog ddhCTP, resulting in enhanced inhibition of viral RNA synthesis and the replication of Crimean-Congo hemorrhagic fever virus and SARS-CoV-2. Together, these findings demonstrate an ISG15-USP18-viperin axis that can be targeted to boost the metabolic antiviral restriction.

## INTRODUCTION

Type I interferons (IFN-I) are central mediators of innate antiviral immunity that induce a rapid transcriptional program establishing a broad antiviral state. Engagement of the type I interferon receptor (IFNAR) activates the JAK-STAT signaling pathway, leading to the induction of hundreds of interferon-stimulated genes (ISGs) that collectively restrict viral replication across diverse virus families^1^. While this response provides powerful antiviral protection, IFN signaling must be tightly regulated. Excessive magnitude or prolonged duration of IFN responses disrupts cellular homeostasis and can lead to harmful inflammatory and metabolic consequences, underscoring the importance of negative feedback mechanisms that limit IFN signaling once antiviral protection has been achieved^1–3^.

Interferon-stimulated gene 15 (ISG15), a ubiquitin-like protein strongly induced by IFN-I, occupies a distinctive position within the interferon system. ISG15 can be covalently conjugated to host and viral proteins through an enzymatic cascade analogous to ubiquitination, a process termed ISGylation^4^. Early studies in mice established ISG15 and ISGylation as antiviral effectors: *Isg15*-deficient mice display increased susceptibility to viral infections, including influenza A and B viruses and herpes simplex virus type 1, and loss of the ISGylation E1 enzyme *UbE1l* results in a similar phenotype^5,6^. These observations positioned ISG15 as a classical antiviral restriction factor in murine models. In contrast, human genetic studies revealed a fundamentally different biology. Individuals with inherited ISG15 deficiency do not exhibit broad susceptibility to viral infections^7^. Instead, they develop Mendelian susceptibility to mycobacterial disease accompanied by pronounced interferon signatures. This paradox was resolved by the discovery that intracellular free human ISG15 stabilizes USP18, a dominant negative regulator of IFNAR signaling, by preventing its proteasomal degradation by skp2^8^.

USP18 suppresses receptor-proximal JAK-STAT activation, and failure to accumulate USP18 results in prolonged STAT1 and STAT2 phosphorylation and sustained ISG expression^8–10^. In ISG15-deficient human cells, USP18 is transcriptionally induced but fails to persist at the protein level, leading to exaggerated interferon signaling^8^. Importantly, this regulatory mechanism is species-specific. Comparative studies have demonstrated that murine USP18 stability is independent of ISG15, allowing mice to regulate IFN signaling without ISG15-mediated stabilization, whereas human cells rely on ISG15 to maintain USP18-dependent negative feedback^11^. This divergence explains why ISG15 deficiency leads to antiviral susceptibility in mice but interferonopathy-like phenotypes in humans and highlights the need to interpret ISG15 biology within a human-specific regulatory framework^4^.

While exaggerated IFN signaling in human ISG15 deficiency has been well documented at the transcriptional level, its consequences at the protein and effector level remain poorly understood. A recent transcriptomic profiling of IFNα-stimulated fibroblasts from ISG15-deficient individuals revealed a broad and sustained upregulation of ISGs compared with controls, reinforcing the role of ISG15 as a negative regulator of IFN output^12^. However, whether this transcriptional hyperresponsiveness causes a global amplification of the IFN response or reinforces specific antiviral effector pathways with distinct functional consequences, remains to be resolved. Recent work has established that ISGs typically function in combinatorial networks rather than as individual dominant effectors^12,13^. Whether ISG15 deficiency broadly amplifies these networks or creates specific regulatory dependencies, remains unclear. Importantly, antiviral protection ultimately depends on the accumulation and activity of effector proteins and metabolites rather than transcript abundance alone. Thus, it remains unclear whether dysregulated IFN signaling in ISG15-deficient human cells broadly amplifies antiviral networks or instead promotes specific metabolic antiviral programs that directly restrict viral replication.

Here, we sought to define how dysregulated IFN signaling ISG15-deficient human cells reshape antiviral effector programs at the protein and metabolic levels. Using quantitative proteomics, genetic reconstitution, and viral infection models, we identify a signaling-to-metabolism axis in which loss of ISG15 amplifies viperin (RSAD2) expression and promotes accumulation of the chain-terminating antiviral nucleotide analog 3′-deoxy-3′,4′-didehydro-cytidine triphosphate (ddhCTP), resulting in enhanced restriction of viral RNA synthesis.

## RESULTS

### ISG15 deficiency amplifies viperin expression following IFN signaling

To define how loss of ISG15 reshapes the interferon-induced effector landscape, we performed TMT-labeling-based quantitative proteomics in isogenic Huh7 Cas9 control cells with intact ISG15 expression (hereafter referred to as wild-type, WT) and in ISG15 knockout (KO) cells following 24h IFN-β stimulation (Figure 1A). This unbiased proteomic analysis identified distinct sets of IFN-regulated proteins in ISG15-deficient cells compared with WT controls, including several canonical ISGs that were enriched among proteins uniquely upregulated in the KO cells (Figure 1B; Supplementary Figure S1A-E, Supplementary Table S1), consistent with defective negative feedback regulation of IFN signaling, as previously described in ISG15-deficient human cells^8^.

**Figure 1.**
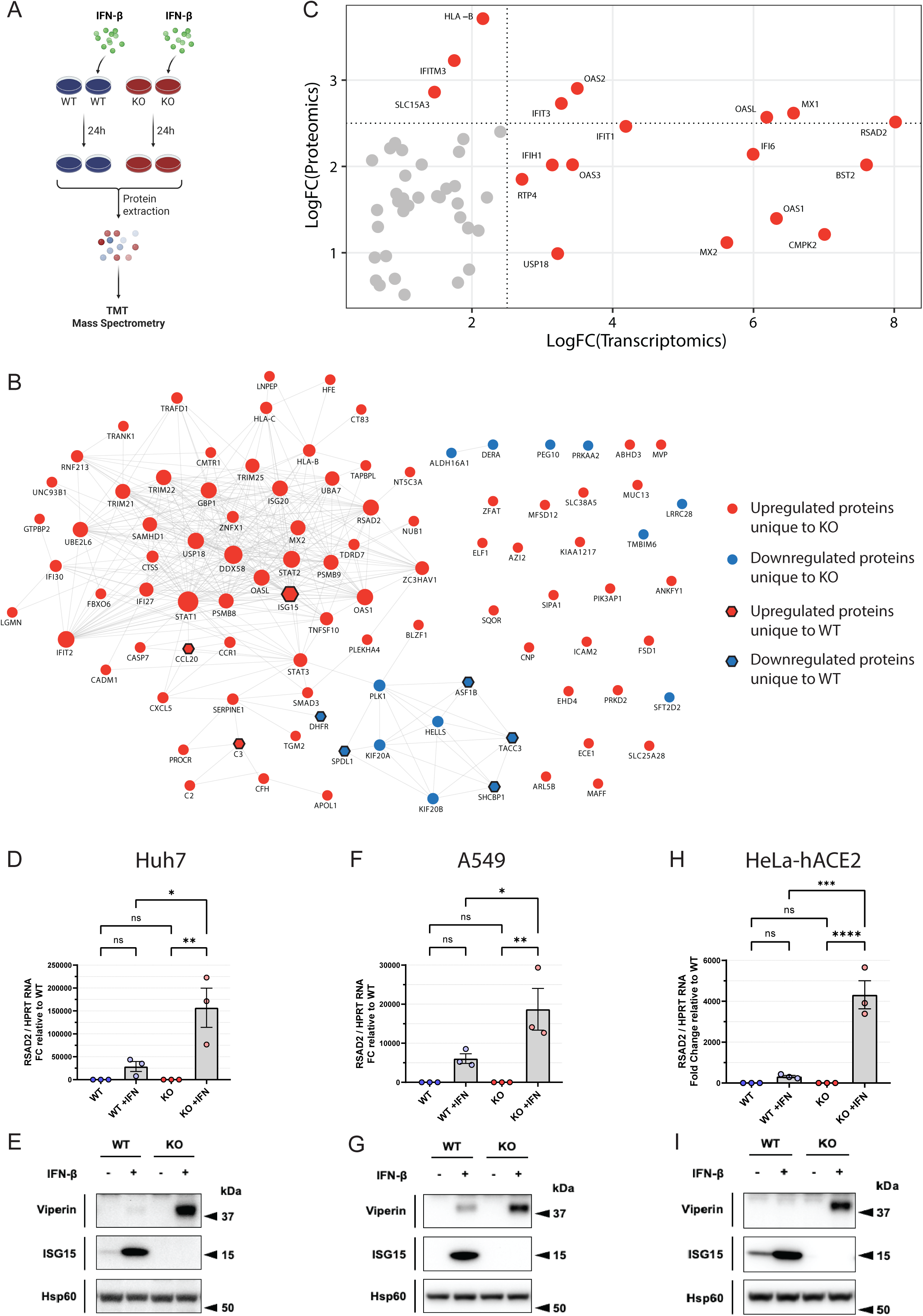
Loss of ISG15 amplifies interferon-driven viperin (RSAD2) induction across epithelial cell types. **(A)** Experimental design for quantitative proteomics. Wild-type (WT) and ISG15 knockout (KO) cells were left untreated or stimulated with IFN-β (1000U/mL) for 24 h prior to protein extraction. Lysates were subjected to tandem mass tag (TMT)-based quantitative mass spectrometry for comparative proteomic profiling. Figure created with BioRender.com. **(B)** Network representation of proteins differentially regulated between WT and ISG15KO cells following IFN-β stimulation. Nodes represent significantly regulated proteins unique to each genotype (adjusted p value as defined in Methods), color-coded as follows: red circles, upregulated proteins unique to ISG15KO; blue circles, downregulated proteins unique to ISG15KO; red hexagons, upregulated proteins unique to WT; blue hexagons, downregulated proteins unique to WT. Node size reflects relative fold-change magnitude. Edges denote experimentally validated or curated functional interactions (STRING-based network integration). **(C)** Association of transcriptomic changes in ISG15-deficient patient derived fibroblasts after IFN-a stimulation^12^ and proteomic responses ISG15 KO Huh7 cells after IFN-β stimulation (This study). Scatter plot showing log2 fold-change (LogFC) at the transcript level (x axis) versus proteome level (y axis). Selected antiviral ISGs are annotated. Dashed lines indicate threshold cutoffs for significant differential expression (as defined in Methods). **(D, F, H)**Quantitative RT-PCR analysis of RSAD2 mRNA expression in WT and ISG15KO cells ± IFN-β (24 h) in three epithelial cell models: **(D)** Huh7, **(F)** A549, and **(H)** HeLa-hACE2. RSAD2 expression was normalized to HPRT and presented as fold-change relative to untreated WT controls. Data are shown as mean ± SEM from independent biological replicates (n ≥ 3). Statistical analysis was performed using ordinary one-way ANOVA; ns not significant; *p < 0.05; **p < 0.01; ***p < 0.001; ****p < 0.0001. **(E, G, I)** Immunoblot validation of viperin (RSAD2) protein expression in **(E)** Huh7, **(G)** A549, and **(I)** HeLa-hACE2 cells ± IFN-β (24 h). ISG15 expression confirms genotype and IFN responsiveness. Hsp60 serves as a loading control. Immunoblots are representative of independent experiments (n ≥ 3).

To place our findings in the context of human ISG15 deficiency, we integrated our proteomic dataset with recent transcriptomic data from IFNα2b-stimulated fibroblasts derived from ISG15-deficient patients^12^. While the patient study defined transcriptional changes associated with exaggerated IFN signaling, it remained unclear how these alterations translate to the protein level. Although our experiments employed IFN-β stimulation while the patient dataset used IFNα2b, both cytokines signal through the shared IFNAR receptor and induce highly overlapping interferon-stimulated gene (ISG) programs. Cross-platform comparison revealed substantial concordance between the datasets, with several canonical ISGs elevated in both systems (Figure 1C; Supplementary Figure S1F). Among these, RSAD2 (viperin) emerged as one of the most consistently and robustly elevated ISGs across the datasets. Given its established role in antiviral nucleotide metabolism, this observation prompted further investigation of viperin regulation and function in the context of ISG15 deficiency.

We next validated viperin induction across multiple epithelial cell models relevant to viral infection. IFN-β stimulation resulted in significantly increased viperin expression at both the mRNA and protein levels in ISG15 KO Huh7 cells compared with WT counterparts (Figure 1D-E). This phenotype was recapitulated in A549 lung epithelial cells and HeLa-hACE2 cells (Figure 1F-I). The consistency across distinct cellular backgrounds indicated that the prominent amplification of viperin among multiple induced ISGs is a general feature of ISG15 loss.

### Loss of ISG15 destabilizes USP18 and enhances early JAK-STAT signaling

The broad enhanced induction of ISGs observed in ISG15-deficient cells compared to cells expressing ISG15, suggests impaired negative feedback regulation of IFN signaling. USP18 is the principal negative regulator of type I IFN signaling, and its deficiency results in IFN hypersensitivity and interferonopathy^10^. To determine whether the differences in IFN signaling account for the increased ISG induction observed in ISG15-deficient cells, we performed a detailed time-course analysis of IFN-β signaling in Huh7 WT and ISG15 KO cells. Following IFN-β stimulation over 24 hours, ISG15-deficient cells displayed increased phosphorylation of STAT1 and STAT2 at early time points compared with WT cells (Figure 2A; Supplementary Figure S2A-D). This enhanced proximal signaling was accompanied by increased induction of downstream ISGs, including viperin, over the same time course (Figure 2A; Supplementary Figure S2E-L), consistent with prior observations in ISG15-deficient patient-derived cells^8^.

**Figure 2.**
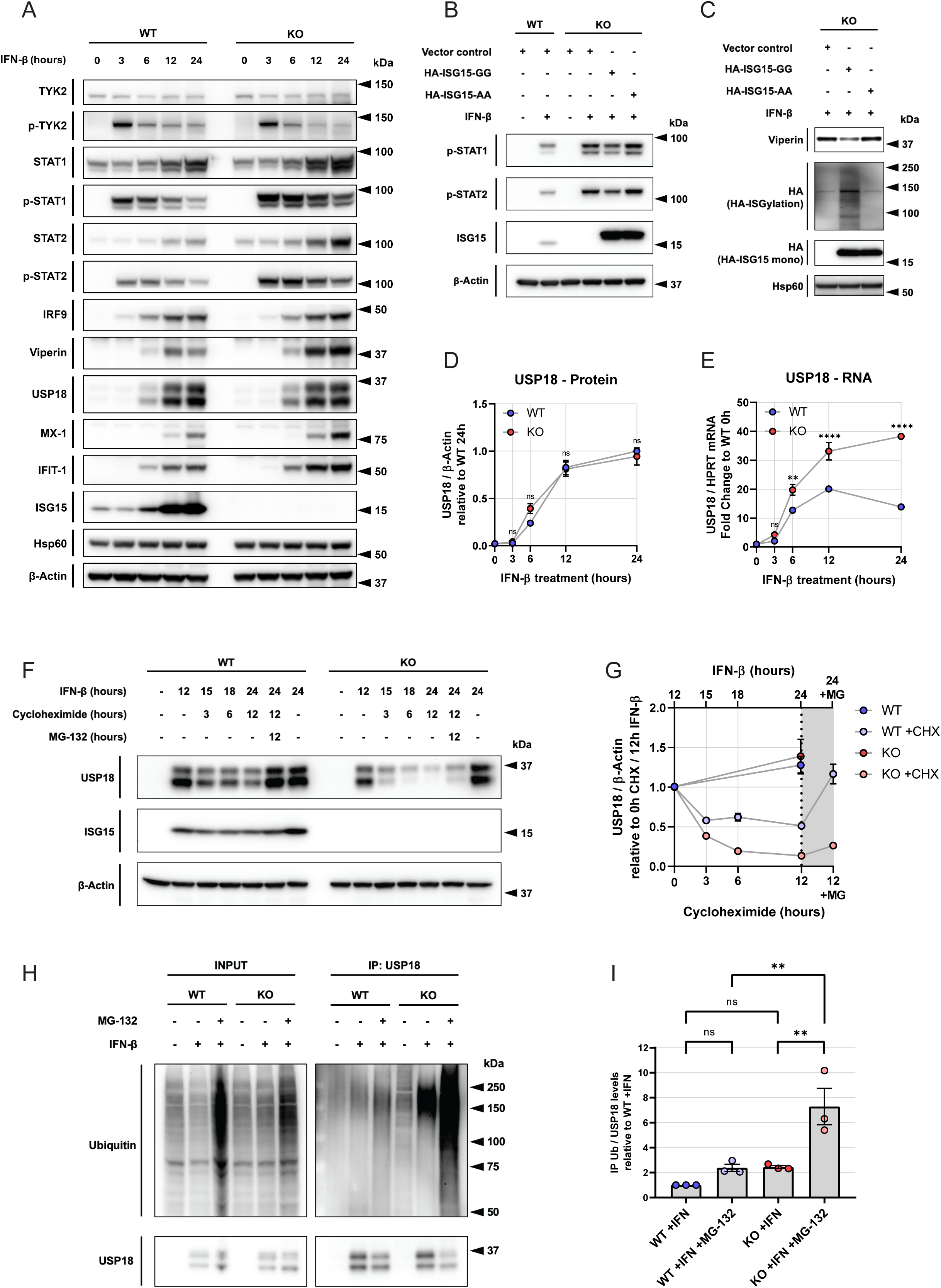
ISG15 stabilizes USP18 and restrains sustained JAK–STAT signaling through protection from proteasomal degradation. **(A)** Time-resolved analysis of IFN-β signaling in WT and ISG15KO cells. Immunoblot of TYK2, phospho-TYK2 (p-TYK2), STAT1, phospho-STAT1 (p-STAT1), STAT2, phospho-STAT2 (p-STAT2), IRF9, Viperin (RSAD2), USP18, MX1, IFIT1, and ISG15 following IFN-β (1000U/mL) stimulation (0-24 h). β-Actin and Hsp60 serve as loading controls. Immunoblots are representative of independent experiments (n ≥ 3) **(B)** Rescue of IFN signaling control by conjugation-competent ISG15. WT and ISG15KO cells were transfected with HA-tagged pcDNA3.1 empty vector control, HA-ISG15-GG (conjugation-competent), or HA-ISG15-AA (conjugation-defective) for 24h, followed by IFN-β (1000U/mL) stimulation for 24h. Immunoblot analysis of p-STAT1, p-STAT2, ISG15, and β-actin as loading control. Immunoblots are representative of independent experiments (n ≥ 3) **(C)** Impact of ISG15 conjugation on viperin expression. KO cells expressing HA-ISG15-GG or HA-ISG15-AA were stimulated with IFN-β (1000U/mL) for 24h and analyzed for viperin protein levels. High-molecular-weight HA-reactive smears indicate global ISGylation in HA-ISG15-GG-expressing cells but not with HA-ISG15-AA. Hsp60 serves as a loading control. Immunoblots are representative of independent experiments (n ≥ 3) **(D)** Quantification of USP18 protein kinetics following IFN-β treatment (0-24 h). USP18 levels were normalized to β-actin and expressed relative to WT 24 h. Data represent mean ± SEM from independent biological replicates (n ≥ 3). Statistical analysis was performed using two-way ANOVA with multiple comparisons; ns, not significant. **(E)** USP18 mRNA induction in WT and ISG15KO cells following IFN-β stimulation (0-24 h). USP18 transcripts were normalized to HPRT and expressed as fold-change relative to WT 0 h. Data represent mean ± SEM (n ≥ 3). Statistical analysis was performed using two-way ANOVA with multiple comparisons; ns not significant; ***p < 0.001; ****p < 0.0001. **(F)** Cycloheximide (CHX) chase assay assessing USP18 protein stability. WT and ISG15KO cells were pretreated with IFN-β (12 h), followed by CHX treatment (50 µg/mL) for 3h, 6h and 12h to inhibit de novo protein synthesis, in the presence or absence of the proteasome inhibitor MG-132 (5µM) for 12h. Immunoblot of USP18. ISG15 expression confirms genotype and IFN responsiveness. β-actin serves as a loading control. Immunoblots are representative of independent experiments (n ≥ 3) **(G)** Quantification of USP18 stability during CHX chase. USP18 levels were normalized to β-actin and expressed relative to 0 h CHX following 12 h IFN-β priming. Data represent mean ± SEM from independent experiments (n ≥ 3). **(H)** Ubiquitination of USP18 in WT and ISG15KO cells. Following IFN-β stimulation (1000U/mL) for 24h and ± MG-132 (5µM) for last 12h, USP18 was immunoprecipitated and probed for ubiquitin by immunoblot. Immunoblot of USP18 serves as a control. Input and IP fractions are shown. Immunoblots are representative of independent experiments (n ≥ 3) **(I)** Quantification of ubiquitinated USP18 species from panel (H). Ubiquitin signal intensity in USP18 IPs was normalized to total USP18 levels and expressed relative to WT + IFN. Data are shown as mean ± SEM (n ≥ 3 independent experiments). Statistical analysis was performed using Ordinary one-way ANOVA; ns not significant; **p < 0.01.

To test whether ISGylation is required to dampen early JAK-STAT activation, we reconstituted ISG15 KO cells with either HA-tagged conjugation-competent ISG15 (HA-ISG15-GG) or a non-conjugatable mutant lacking the C-terminal di-glycine motif (HA-ISG15-AA). Ectopic expression of HA-ISG15-GG suppressed elevated STAT1 and STAT2 phosphorylation, whereas HA-ISG15-AA failed to do so (Figure 2B; Supplementary Figure S2M-N). In parallel, the expression of ISG15-GG, but not ISG15-AA, attenuated IFNβ-induced viperin expression in KO cells (Figure 2C; Supplementary Figure S2O). These findings indicate that conjugation-competent ISG15 is required to restrain IFN signaling and downstream viperin induction, consistent with recent evidence that ISG15 conjugation contributes to IFN antiviral activity in human cells^14^.

We next investigated whether ISG15 regulates USP18 protein stability during IFN signaling. Quantification of the Figure 2A immunoblots revealed that USP18 protein accumulated with similar kinetics in WT and ISG15 KO cells following IFN-β stimulation (Figure 2D), despite significantly higher USP18 mRNA levels in KO cells (Figure 2E). This discrepancy suggested post-transcriptional regulation of USP18 in the absence of ISG15. To directly test USP18 stability, we performed cycloheximide (CHX) chase experiments, in which inhibition of protein synthesis allows measurement of protein degradation over time. CHX chase experiments revealed accelerated USP18 turn-over in ISG15-deficient cells (Figure 2F-G). Immunoprecipitation further demonstrated increased poly-ubiquitination of USP18 in ISG15 KO cells, visible as a high-molecular-weight ubiquitin smear in USP18 immunoprecipitates and enhanced response to the proteasome inhibitor MG132 (Figure 2H-I), consistent with increased proteasomal targeting as previously reported^8,15^. Together, these data indicate that loss of ISG15 promotes ubiquitin-mediated turnover of USP18, thereby contributing to exaggerated early JAK-STAT activation and enhanced induction of downstream ISGs, including viperin.

### Loss of ISG15 promotes viperin-dependent ddhCTP accumulation

Given the amplified induction of ISGs in ISG15-deficient cells upon IFN-β treatment, we next asked whether this dysregulated IFN state translated into measurable antiviral effects. To this end, we examined whether this enhanced antiviral state would restrict replication of Crimean-Congo hemorrhagic fever virus (CCHFV), a negative-sense RNA virus of the family Nairoviridae encoding Ub and ISG15 specific protease that circumvent IFN induced antiviral state^16^. During experiments in which Huh7 cells were transfected with an empty vector prior to infection, ISG15 KO cells displayed significantly lower CCHFV RNA levels compared with WT cells (Supplementary Figure S3A). Consistent with enhanced interferon signaling in ISG15-deficient cells, RSAD2 transcript levels were elevated under these conditions (Supplementary Figure S3B). Using a CCHFV/ZsGreen (ZsG) reporter virus, we observed a similar phenotype in A549 cells,where an ISG15 deficiency similarly resulted in reduced ZsG reporter expression over time (Supplementary Figure S3C). These findings revealed a reproducible antiviral phenotype associated with ISG15 loss, consistent with the amplified ISG network we observed.

Because RSAD2 (viperin) is a radical S-adenosylmethionine enzyme that catalyzes the conversion of cytidine triphosphate (CTP) into the antiviral nucleotide analog ddhCTP^17^, we asked whether the antiviral phenotype observed in ISG15-deficient cells could be explained by viperin-dependent remodeling of nucleotide metabolism. ddhCTP has been shown to interfere with viral RNA-dependent RNA polymerases by acting as an inhibitory or chain-terminating substrate and, under certain conditions, by impairing viral RNA translation^17,18^. We therefore tested whether ISG15 loss promotes accumulation of ddhCTP following IFN stimulation.

A schematic overview of the experimental workflow is shown in Figure 3A. We first quantified canonical ribonucleoside triphosphates (rNTPs) using an RNA polymerase-based enzymatic assay^19^. ATP and GTP levels were comparable between ISG15-intact control (WT) and ISG15 KO cells and were not significantly altered by IFN-β treatment (Figure 3B). In contrast, UTP and CTP signals were reduced in ISG15 KO cells at baseline, with a further decrease in apparent CTP levels following IFN-β stimulation (Figure 3B).

**Figure 3.**
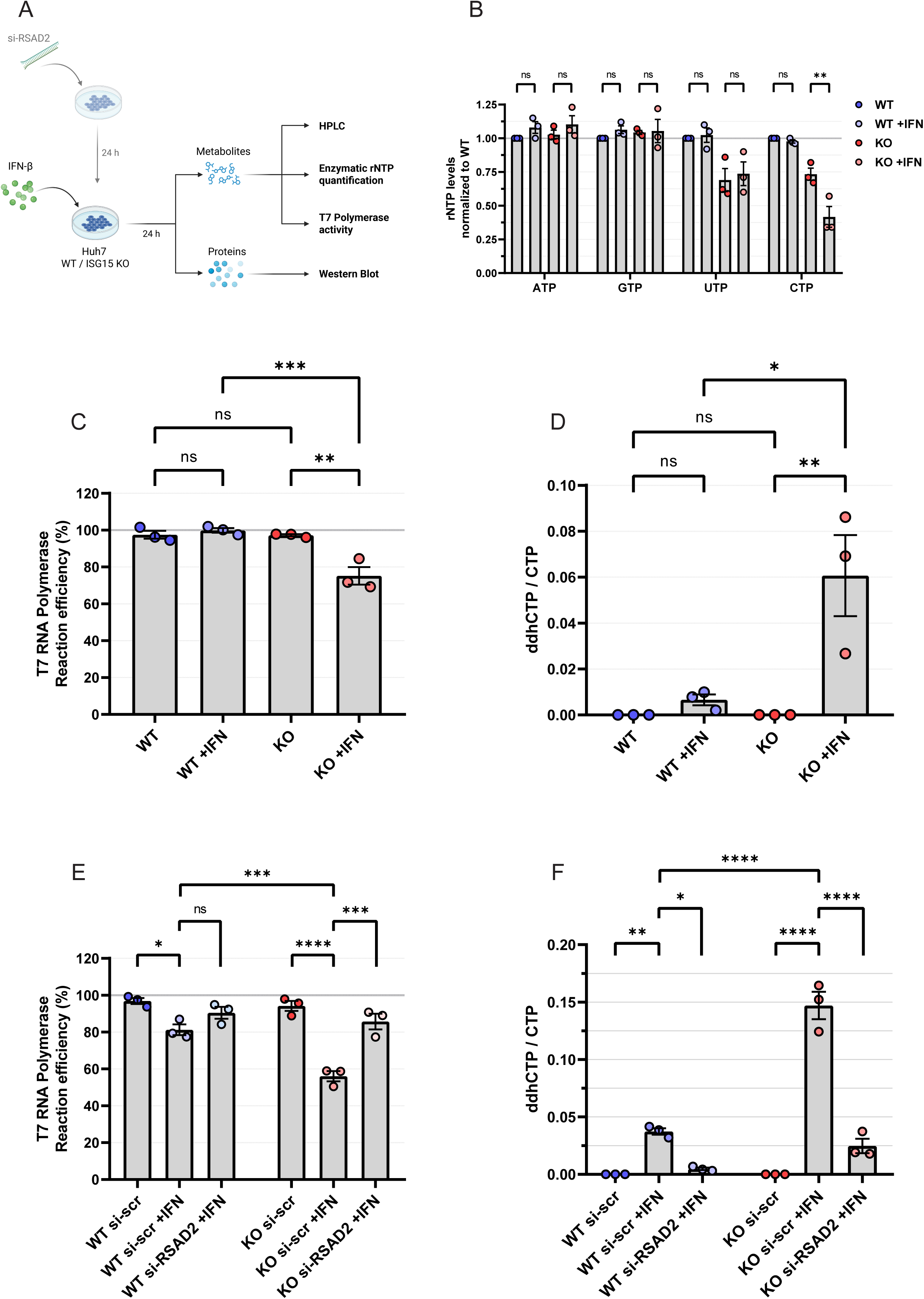
ISG15 deficiency amplifies viperin-dependent ddhCTP production and suppresses RNA polymerase activity without global nucleotide depletion. **(A)** Experimental workflow. Huh7 WT and ISG15KO cells were transfected with siRNA targeting RSAD2 (viperin) or scrambled control (si-scr), followed by IFN-β stimulation (24 h). Cells were harvested for (i) metabolite extraction and HPLC-based ddhCTP quantification, (ii) enzymatic rNTP quantification assays, (iii) in vitro T7 RNA polymerase activity assays, and (iv) immunoblot validation. Figure created with BioRender.com. **(B)** Quantification of intracellular ribonucleotide triphosphate (rNTP) levels (ATP, GTP, UTP, CTP) in WT and ISG15KO cells ± IFN-β (24 h) using an enzymatic microplate assay based on fluorogenic *in vitro* transcription. rNTP levels were normalized to untreated WT controls. Data represent mean ± SEM from independent biological replicates (n ≥ 3). Statistical analysis was performed using two-way ANOVA with multiple comparisons; ns not significant; **p < 0.01. **(C)** T7 RNA polymerase interference test using total cellular metabolite extracts from WT and ISG15KO cells ± IFN-β (24 h). The Standard Addition Method was utilized to quantify the interference in CTP quantifications utilizing T7 RNA polymerase. T7 RNA polymerase reaction efficiency is expressed as percentage relative to WT control. Data represent mean ± SEM from independent experiments (n ≥ 3). Statistical analysis was performed using ordinary one-way ANOVA; ns not significant; **p < 0.01; ***p < 0.001. **(D)** Intracellular ddhCTP / CTP ratio quantified by HPLC in WT and ISG15KO cells ± IFN-β (24 h). Data represent mean ± SEM from independent experiments (n ≥ 3). Statistical analysis was performed using ordinary one-way ANOVA; ns not significant; *p < 0.05. **(E)** T7 RNA polymerase activity following RSAD2 knockdown. WT and ISG15KO cells were treated with si-scr or si-RSAD2 and stimulated with IFN-β (1000U/mL) for 24 h. The Standard Addition Method was used to quantify the interference in CTP quantification of cellular metabolite extracts utilizing T7 RNA polymerase. T7 RNA polymerase reaction efficiency is expressed as percentage relative to WT si-scr control. Data represent mean ± SEM from independent experiments (n ≥ 3). Statistical analysis was performed using ordinary one-way ANOVA; *p < 0.05; ***p < 0.001; ****p < 0.0001. **(F)** ddhCTP / CTP ratio following RSAD2 knockdown. WT and ISG15KO cells were treated with si-scr or si-RSAD2 and stimulated with IFN-β (1000U/mL) for 24 h. The intracellular ddhCTP / CTP ratio was quantified by HPLC. Data are shown as mean ± SEM (n ≥ 3). Statistical analysis was performed using ordinary one-way ANOVA; ***p < 0.001; ****p < 0.0001.

Since this assay relies on bacteriophage T7 (or SP6) RNA polymerase to extend RNA under limiting nucleotide conditions, we considered whether the apparent reduction in CTP reflected true depletion or functional interference by a CTP analog. To directly test this possibility, we repurposed the system as a T7 RNA polymerase interference assay. Metabolite extracts from IFN-treated ISG15 KO cells inhibited synthesis of full-length (102 nt) transcripts under limiting CTP conditions, whereas extracts from WT cells did not (Figure 3C). This indicated the presence of a CTP-competitive inhibitory species in ISG15-deficient samples.

To quantify nucleotide species directly and independently of RNA polymerase activity, we performed high-performance liquid chromatography (HPLC). Importantly, HPLC-based measurements of ATP, GTP, and UTP strongly correlated with the enzymatic assay (Supplementary Figure S4A), validating the overall accuracy of nucleotide quantification. However, in contrast to the enzymatic readout, HPLC analysis did not detect a reduction in absolute CTP levels in IFN-treated ISG15 KO cells (Supplementary Figure S4A). When CTP and ddhCTP were resolved and quantified together (Supplementary Figure S4B), a marked increase in ddhCTP was observed specifically in IFN-treated ISG15 KO cells, whereas ddhCTP remained low or undetectable in WT cells (Figure 3D). These results demonstrate that the apparent CTP reduction detected by the enzymatic assay reflects ddhCTP-mediated interference rather than depletion of endogenous CTP pools.

To determine whether ddhCTP accumulation depended on viperin activity, we depleted RSAD2 using siRNA and confirmed efficient knockdown by immunoblotting (Supplementary Figure S4C). RSAD2 knockdown abolished the inhibitory effect of ISG15 KO metabolite extracts in the T7 polymerase assay (Figure 3E), indicating that the interfering effect was viperin-dependent. Consistently, HPLC analysis demonstrated a marked reduction in ddhCTP levels upon RSAD2 depletion (Figure 3F, Supplementary Figure S4D). Also, enzymatic CTP measurements in RSAD2-depleted cells no longer showed the apparent reduction observed in control ISG15 KO cells (Supplementary Figure S4E), and HPLC confirmed that absolute CTP levels remained unchanged (Supplementary Figure S4D).

Together, these data demonstrate that loss of ISG15 promotes viperin-dependent accumulation of ddhCTP following IFN stimulation, establishing a direct biochemical link between dysregulated IFN signaling in ISG15-deficient cells and accumulation of a metabolite capable of restricting viral RNA polymerase activity.

### Viperin-dependent ddhCTP accumulation contributes to viral restriction in ISG15-deficient cells

To directly assess the impact of viperin-derived ddhCTP on viral RNA synthesis, we monitored CCHFV/ZsG replication in A549 cells, using ZsG fluorescence as a readout for viral nucleoprotein expression and replication kinetics. Depletion of RSAD2 restored reporter virus signal in ISG15 KO cells to the levels observed in WT cells (Figure 4A). Furthermore, the virus-ZsG levels negatively correlated with viperin mRNA levels, supporting a direct link between enhanced viperin expression and reduced viral replication (Figure 4B). Infectious virus production measured by TCID₅₀ was similarly reduced in ISG15 KO compared to WT cells (Supplementary Figure S5A). RSAD2 depletion significantly restored CCHFV/ZsG infectious titers, confirming that viperin contributes to the enhanced viral restriction observed in ISG15-deficient cells, even in the absence of exogenous IFN treatment. While these data support a role for viperin in restricting CCHFV replication, our experiments did not distinguish between inhibitory effects on viral entry, replication, or assembly. To isolate effects on viral RNA synthesis from entry or assembly, we employed a minigenome system that reports CCHFV L-RNA-dependent RNA polymerase activity^20^ (Figure 4C; Supplementary Figure S5B-C). To avoid confounding effects from T7 RNA polymerase-dependent transcription required for reporter expression, we first expressed the viral N and L proteins, followed by IFN-β treatment, and subsequently transfected *in vitro*-transcribed nano-luciferase-expressing minigenome RNA. Under these conditions, IFNβ-treated ISG15 KO cells exhibited a marked reduction in minigenome-driven nano-luciferase activity compared with WT cells, indicative of impaired L polymerase-dependent viral RNA synthesis (Figure 4D). Notably, knockdown of RSAD2 in ISG15 KO cells rescued minigenome activity, restoring luciferase signals to near levels observed in WT cells (Figure 4E). These results further imply viperin as a dominant restriction factor in ISG15-deficient cells.

**Figure 4.**
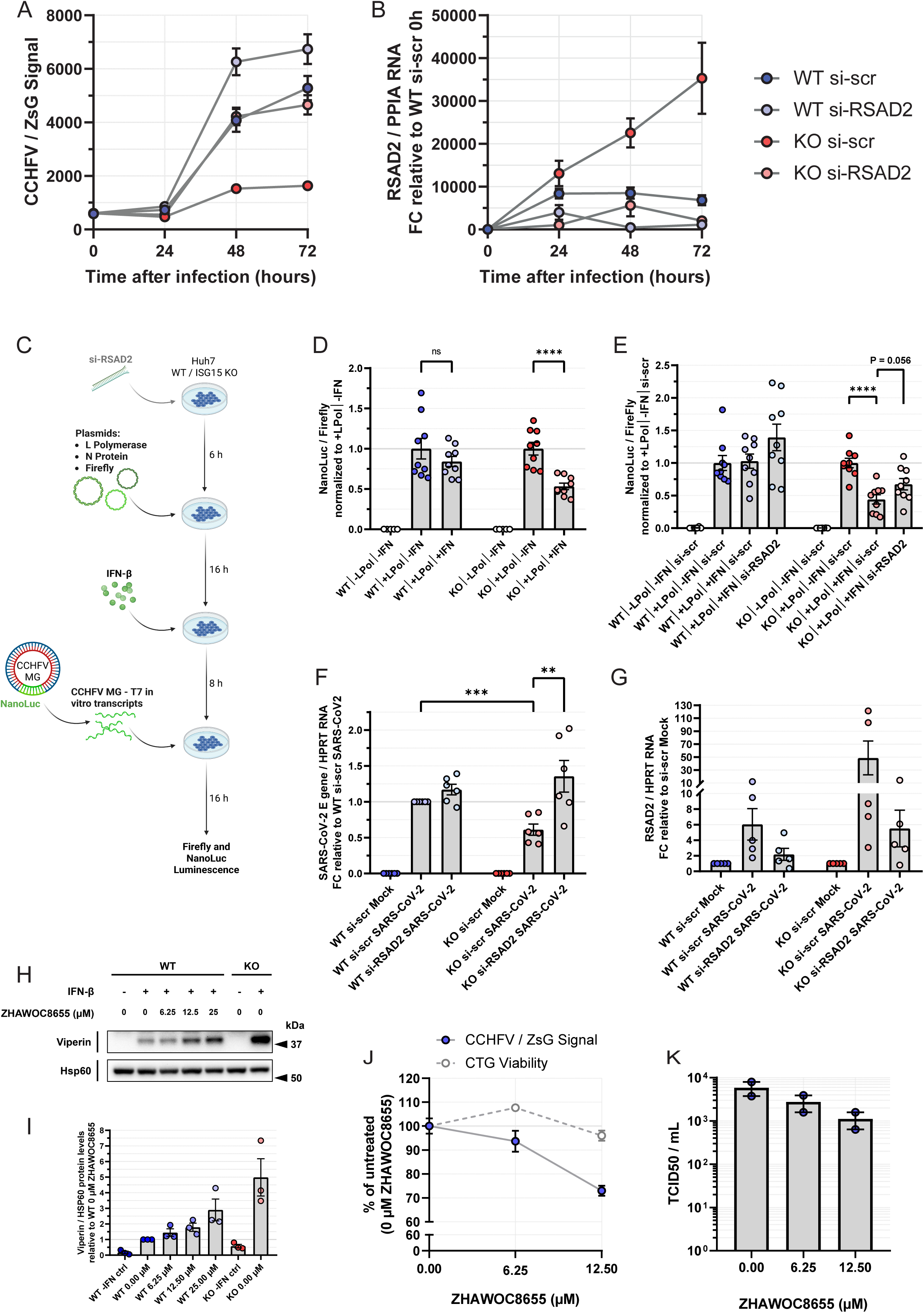
ISG15 constrains viperin-dependent antiviral restriction of viral polymerases in SARS-CoV-2 and CCHFV infection models. **(A)** Replication kinetics of CCHFV/ZsG reporter virus in WT and ISG15KO cells ± RSAD2 knockdown. Cells were transfected with si-scr or si-RSAD2 and infected with recombinant CCHFV expressing ZsGreen. Viral replication was monitored by ZsG fluorescence at 0, 24h, 48h and 72 h post-infection. Data are shown as mean ± SEM from 2 independent experiments each having 4 technical replicates. **(B)** RSAD2 mRNA induction during CCHFV infection. Quantitative RT-PCR analysis of RSAD2 transcripts normalized to PPIA and expressed relative to WT si-scr at 0 h. Data are shown as mean ± SEM from 2 independent experiments each having 3 technical replicates. **(C)** Schematic of the CCHFV minigenome (MG) assay workflow. WT and ISG15KO Huh7 cells were transfected with siRNA (si-scr or si-RSAD2), followed by transfection of plasmids encoding CCHFV L polymerase, N protein, and Firefly luciferase reporter. IFN-β was added prior to transfection of in vitro–transcribed T7-driven NanoLuc CCHFV Minigenome RNA. Polymerase activity was quantified as NanoLuc/Firefly luminescence ratio. Figure created with BioRender.com. **(D)** CCHFV minigenome activity ± IFN-β (1000U/mL). L Polymerase-dependent NanoLuc signal was normalized to Firefly luciferase and expressed relative to +LPol/–IFN condition. Data represent mean ± SEM from 3 biological replicates each having independent technical replicates. Statistical analysis was performed using unpaired t-test; ns not significant; ****p < 0.0001. **(E)** Effect of RSAD2 knockdown on CCHFV minigenome suppression. WT and ISG15KO cells were transfected with si-scr or si-RSAD2 followed by plasmid transfection and CCHFV MG RNA transfection ± IFN-β (1000U/mL). L Polymerase-dependent NanoLuc signal was normalized to Firefly luciferase and expressed relative to +LPol/–IFN/si-scr condition. Data represent mean ± SEM from 3 biological replicates each having independent technical replicates. Statistical analysis was performed using unpaired t-test; ****p < 0.0001. **(F)** SARS-CoV-2 infection in WT and ISG15KO cells ± RSAD2 knockdown. Viral replication was assessed by E gene RNA quantification normalized to HPRT and expressed relative to WT si-scr SARS-CoV-2 condition. Data represent mean ± SEM from 3 biological replicates each having technical duplicates. Statistical analysis was performed using unpaired t-test; ns not significant; **p<0.01; ***p<0.001 **(G)** RSAD2 mRNA levels during SARS-CoV-2 infection. RSAD2 mRNA quantification normalized to HPRT and expressed relative to si-scr mock. Data represent mean ± SEM from 3 biological replicates with either singlet or technical duplicate. **(H)** Immunoblot analysis of viperin protein levels following treatment with the USP18-ISG15 interaction inhibitor ZHAWOC8655 (0-25 µM) ± IFN-β in WT cells. KO cells ± IFN-β was used as a comparative control. Hsp60 serves as loading control. Immunoblots are representative of independent experiments (n ≥ 3) **(I)** Quantification of viperin protein levels from panel (H), normalized to Hsp60 and expressed relative to WT 0 µM ZHAWOC8655. Data are shown as mean ± SEM from independent biological replicates (n ≥ 3). **(J)** Effect of ZHAWOC8655 on CCHFV replication and cell viability. WT and ISG15KO cells were infected with CCHFV/ZsG reporter virus and co-treated with ZHAWOC8655 (0-12.5 µM). CCHFV/ZsG signal and CellTiter-Glo (CTG) viability are expressed as percentage of CCHFV infected / ZHAWOC8655 untreated (0 µM). Data are shown as mean ± SEM from 3 technical replicates. **(K)** Infectious virus titers (TCID50/mL) following ZHAWOC8655 treatment (0-12.5 µM). Data are shown as mean ± SEM from 2 independent biological replicates.

To assess whether this mechanism extends beyond CCHFV, we examined SARS-CoV-2 replication in HeLa-hACE2 cells. Consistent with our observations for CCHFV, SARS-CoV-2 infected ISG15 KO cells also displayed reduced viral RNA levels, as measured by E-gene quantification. Depletion of RSAD2 increased viral RNA abundance in ISG15 KO cells, indicating that viperin similarly contributes to restriction of SARS-CoV-2 replication (Figure 4F-G). While viral RNA polymerases differ in structure and requirements for auxiliary proteins, these data support a model in which elevated viperin expression and ddhCTP accumulation impose a general constraint on RNA virus replication.

### Pharmacologic perturbation of the ISG15-USP18 axis biases viperin induction

Finally, we explored whether pharmacologic perturbation of the ISG15-USP18 axis recapitulates aspects of ISG15 deficiency. A recently described small-molecule inhibitor, ZHAWOC8655, disrupts USP18-ISG15 interactions and inhibits USP18 activity^21^. Treatment with ZHAWOC8655 at non-toxic concentrations (Supplementary Figure S6A) and IFN-b treatment increased viperin protein expression in A549 cells (Figure 4H-I). Notably, other ISGs examined, including MX1, IFIT1, and USP18, were not similarly induced (Supplementary Figure S6B-E).

Functionally, ZHAWOC8655 treatment of A549 WT cells initiated at the time of infection resulted in a dose-dependent reduction of ZsG reporter fluorescence (Figure 4J) and infectious virus production (Figure 4K) following 24 hours of infection. These data indicate that pharmacologic disruption of the ISG15-USP18 axis enhances viperin expression and restricts viral replication, although whether this antiviral effect is solely mediated by viperin remains to be determined.

## DISCUSSION

Type I interferons must deliver rapid and effective antiviral protection while avoiding the tissue damage and metabolic stress associated with prolonged JAK-STAT activation and sustained ISG expression^1^. The central finding of this study is that human ISG15 functions as a homeostatic regulator of IFN-I signaling, and that the loss of this feedback does not simply amplify the interferon response globally but skews the IFN effector landscape toward viperin-driven nucleotide pool remodeling. Using unbiased proteomics, genetic reconstitution, nucleotide quantifications, and infection models, we define a signaling-to-metabolism axis in which defective IFN restraint in ISG15-deficient cells creates a hostile environment for viral RNA synthesis. Within this context, viperin emerges as a prominent downstream effector of dysregulated IFN signaling.

USP18 is a negative regulator of IFN signaling at the receptor-proximal level, and loss of USP18 results in IFN hypersensitivity and inflammatory pathology^10,22^. In ISG15-deficient human cells, USP18 is transcriptionally induced but has high turnover rate at the protein level, leading to prolonged STAT1 and STAT2 phosphorylation and enhanced ISG induction^8,23^. We recapitulate and extend this mechanism in epithelial cell models, demonstrating accelerated USP18 degradation and increased ubiquitination in the absence of ISG15, accompanied by enhanced early JAK-STAT activation. Importantly, restoration of conjugation-competent ISG15 but not a non-conjugatable mutant normalized STAT phosphorylation and suppressed viperin induction, establishing causal links between the ISG15 C-terminal motif and restraint of IFN signaling, and altered ISG output. These findings are consistent with recent mechanistic refinements showing that specific ISG15 C-terminal determinants are required for effective IFN control, beyond simple stabilization of USP18 protein abundance^14,15^.

These findings indicate that loss of ISG15 does not merely amplify interferon signaling but reshapes the downstream effector landscape. Among the ISGs amplified in ISG15-deficient cells, viperin emerged as a key effector because it is not merely a signaling protein but an enzyme that directly modifies nucleotide pools by converting CTP into the antiviral nucleotide analog ddhCTP^17^. A potential translational implication of this finding is targeting the ISG15-USP18-viperin-ddhCTP axis as a broad-spectrum antiviral strategy. ddhCTP interferes with RNA synthesis, a biochemical step shared by many RNA viruses, rather than virus-specific entry or protease functions. Our data extend the antiviral relevance of ddhCTP to CCHFV infection and identify an upstream regulatory pathway controlling its accumulation, with potential for the development of new therapeutic interventions. In this context, a recently described small molecule, ZHAWOC8655, disrupts USP18-ISG15 interactions and inhibits USP18 activity *in vitro*^21^. At sub-toxic concentrations, this compound increased viperin protein abundance without uniformly elevating other ISGs and significantly suppressed CCHFV replication. These observations support the concept that the USP18-ISG15 regulatory checkpoint is chemically addressable and that viperin may represent a particularly potent downstream effector for antiviral interventions. However, genetic and mechanistic studies demonstrate that insufficient USP18-mediated restraint can drive severe interferonopathies^8,10^. Therefore, any therapeutic strategy targeting this pathway must carefully balance antiviral benefit against risks to cellular homeostasis. Such interventions might be particularly viable in the context of acute infections, where shorter treatment windows could minimize adverse effects while maximizing therapeutic benefit.

Beyond antiviral defense, dysregulated viperin activity may also have pathophysiological consequences. For example, inherited interferonopathies linked to ISG15 and USP18 could result in chronic ddhCTP accumulation that could contribute to pathology. ddhCTP is not inherently selective for viral RNA polymerases as mitochondrial RNA polymerase can misincorporate it, leading to premature termination of mitochondrial transcripts^24^. The significance of this host-cell effect is still unclear.

From an evolutionary perspective, our data identify ISGylation as a functionally constrained interface in virus-host interactions. Reconstitution experiments demonstrated that conjugation-competent ISG15 (ISG15-GG), but not the C-terminal di-glycine mutant ISG15-AA, suppressed the elevated viperin induction observed in ISG15-deficient cells following interferon stimulation. Because the di-glycine motif is required for ISG15 conjugation to target proteins, these findings indicate that the integrity of the ISG15 C-terminus is necessary to restrain excessive interferon signaling and downstream ISG induction. Rather than demonstrating a direct requirement for ISGylation, these results suggest that the functional architecture of the ISG15 system, including both free and conjugated forms, contributes to maintaining balanced antiviral responses. This regulatory interface may therefore represent a selectively vulnerable node that viruses repeatedly target during host adaptation.

These observations support a model in which the dual functions of ISG15 create a regulatory axis that can be exploited by viral immune evasion strategies. While free intracellular ISG15 stabilizes USP18 and dampens IFNAR signaling, ISGylation modifies host proteins involved in antiviral effector pathways. Consistent with this framework, diverse RNA viruses, including nairoviruses (e.g., CCHFV) and coronaviruses (e.g., SARS-CoV-2) encode deISGylases that remove ISG15 conjugates from host proteins^16,25^. Reversal of ISGylation by viral deISGylases would be expected to attenuate ISGylation-dependent antiviral effector mechanisms while leaving intact the pool of free ISG15 that stabilizes USP18 and suppresses IFN signaling. Within the ISG15-USP18 negative feedback loop controlling IFNAR signaling, this shift could favor dampening of antiviral effector pathways without provoking excessive interferon responses that would otherwise limit viral replication. The relevant substrates and temporal dynamics of this regulatory axis remain to be defined.

This interpretation is consistent with previous observations that ISG15 is a frequent target of viral immune evasion and also highlights the importance of species-specific ISG15 biology. In murine systems, ISG15/ISGylation can directly restrict viruses through covalent modification of viral or host proteins, such as ISGylation of influenza A virus NS1^26^. In human cells, ISG15 also exerts direct antiviral effects at distinct stages, including inhibition of HIV-1 and Ebola virus budding through interference with ESCRT-dependent processes^27,28^. Many viruses counteract ISG15 by encoding deISGylases, underscoring the selective pressure exerted by this pathway^4^. Our findings extend this framework by linking dysregulated interferon signaling in ISG15-deficient cells to enhanced induction of the antiviral effector viperin and accumulation of the antiviral nucleotide analog ddhCTP. For nairoviruses, this is particularly relevant, as CCHFV encodes an OTU-domain protease with deubiquitinase and deISGylase activity that broadly antagonizes ubiquitin- and ISG15-dependent innate immune signaling^16^. Similar strategies are employed by unrelated RNA viruses, including coronaviruses, whose papain-like proteases exhibit potent deISGylase activity toward human ISG15^25,29^. Together, these observations highlight the ISG15 pathway as a central point of evolutionary conflict between host antiviral defense mechanisms and viral immune evasion strategies.

Some limitations of this study should be acknowledged. The relationship between ddhCTP-to-CTP ratio and the potency against diverse RNA polymerase families remains to be defined. Although the GG-versus-AA rescue highlights the importance of the ISG15 C-terminus, the precise contributions of covalent ISGylation versus noncovalent interactions will benefit from future separation-of-function analyses. Finally, while the inhibitor data provide a proof-of-principle, direct demonstration that pharmacologic perturbation recapitulates the biochemical signatures of genetic ISG15 loss remains an important next step.

In summary, this study refines the mechanism by which ISG15 maintains IFN homeostasis through stabilization of USP18 in humans. When this checkpoint is removed, exaggerated IFN signaling robustly enhances viperin expression, leading to unrestrained ddhCTP accumulation and potent restriction of viral RNA synthesis. Our findings urge further studies to delineate the therapeutic potential of targeting this regulatory pathway and also examine the role of ddhCTP in relevant interferonopathies.

## METHODS

### Key resources table

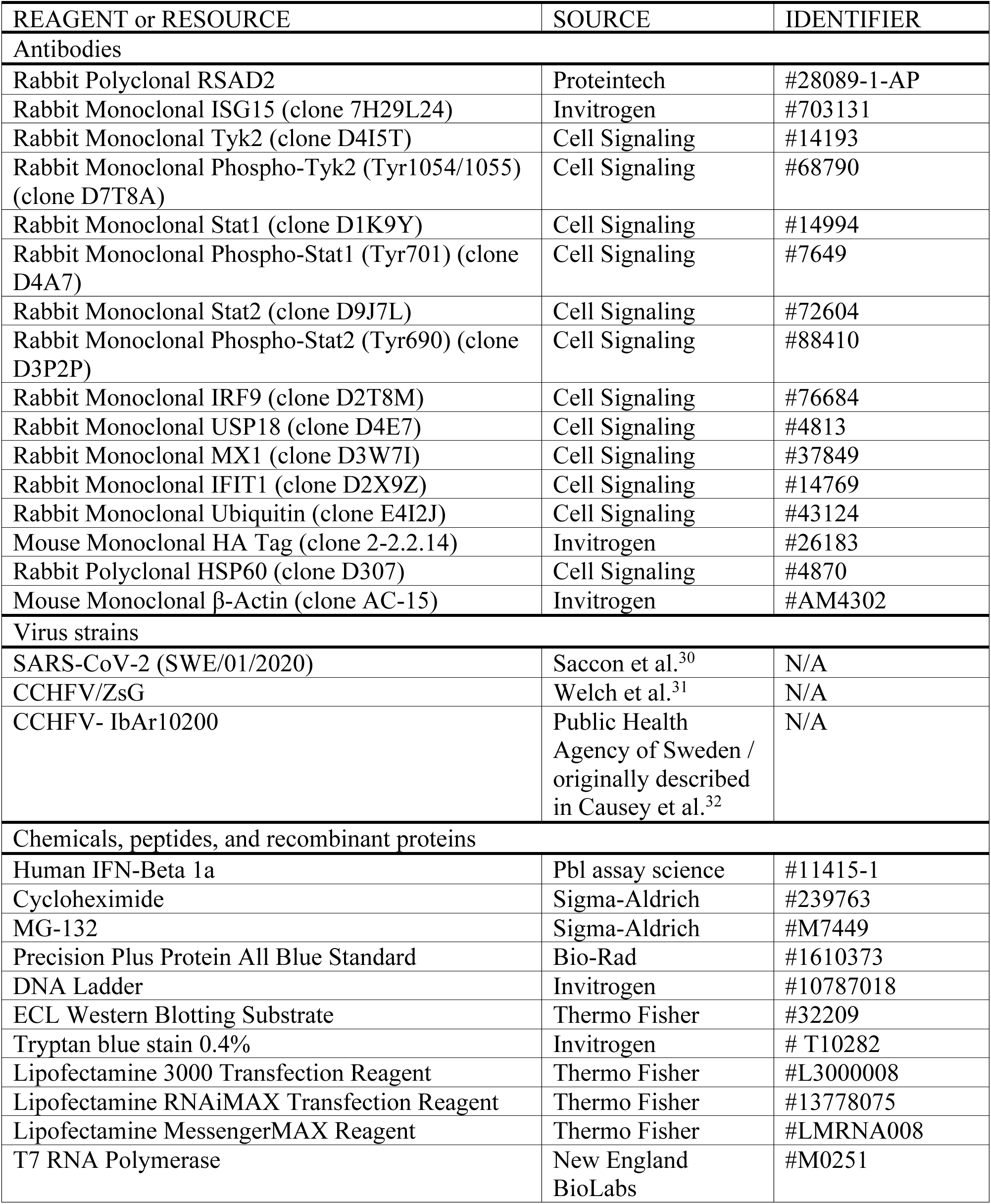
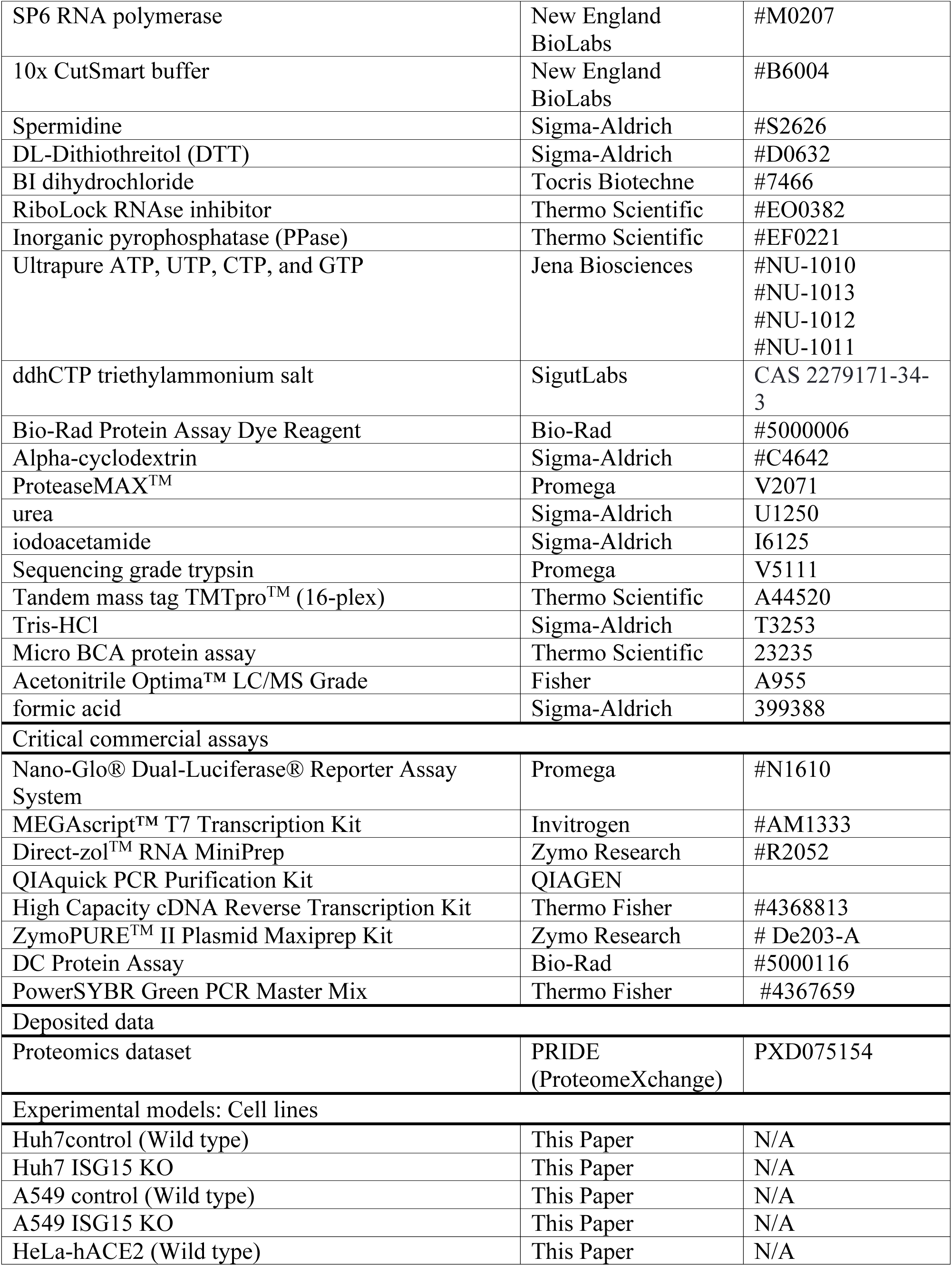
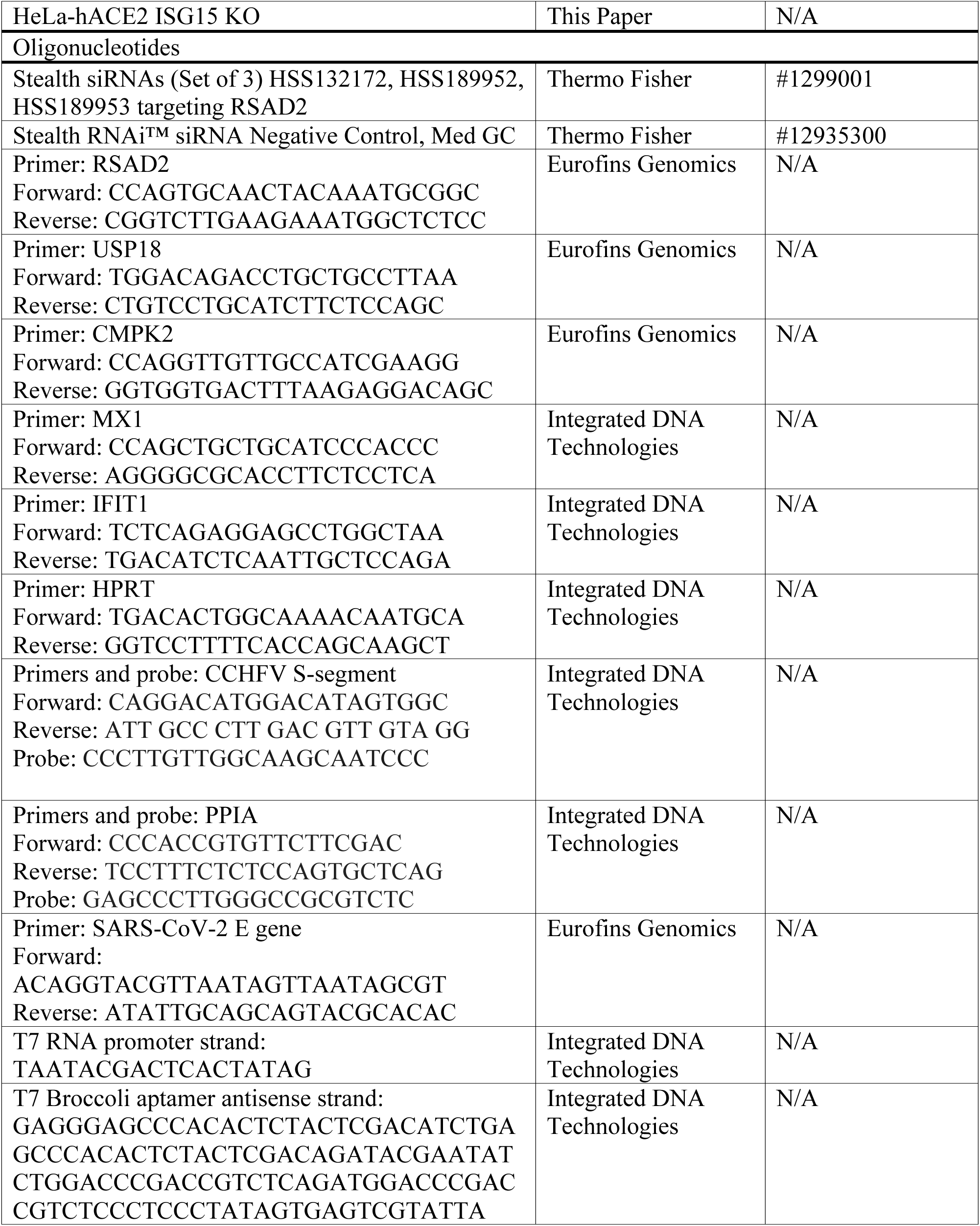
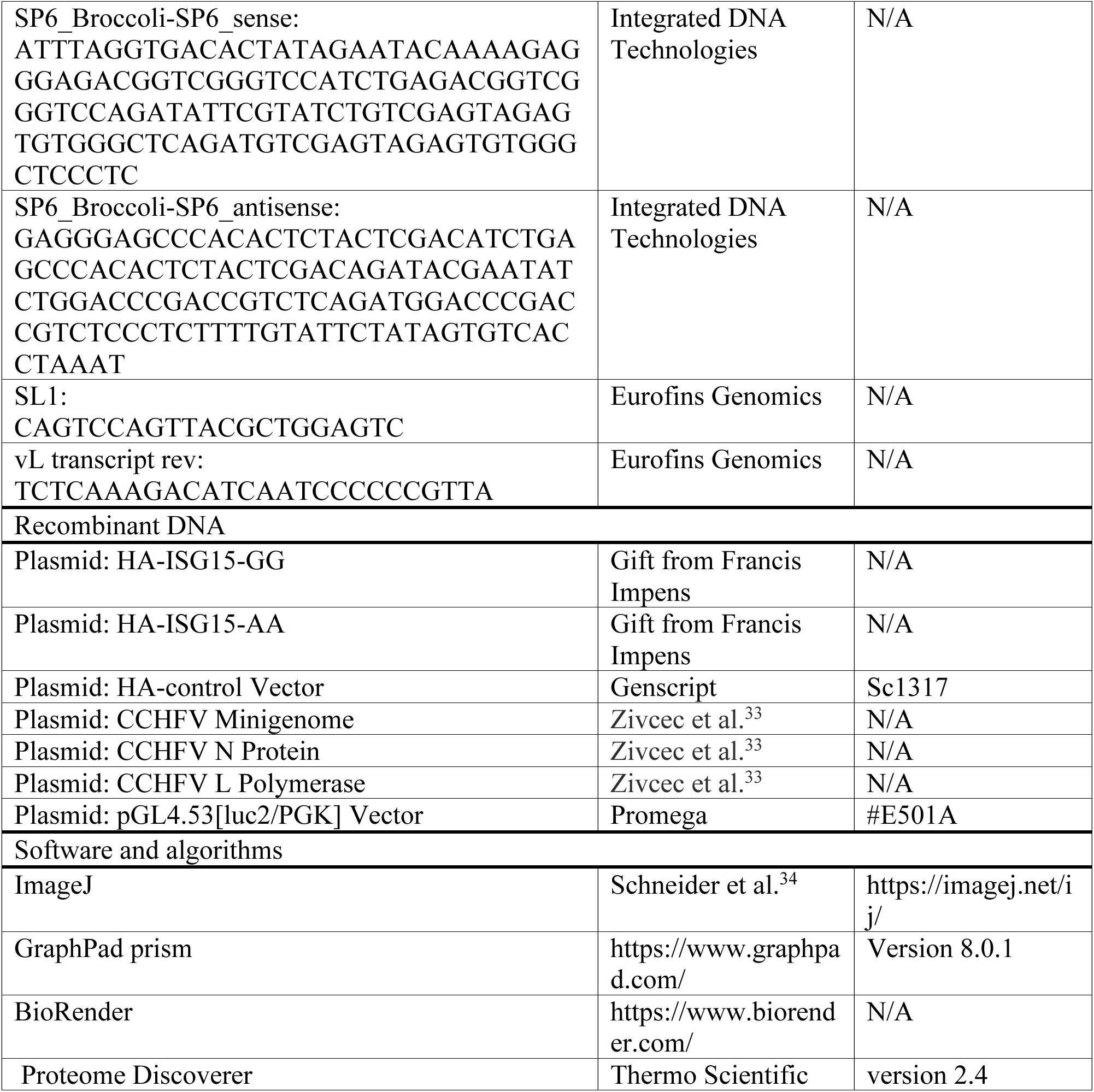

### Experimental Model and Cell Lines

All cell lines were maintained in high-glucose DMEM (Sigma; #D6429-500ML) supplemented with 10% fetal bovine serum (FBS; Gibco; #10500064), 1x Antibiotic-Antimycotic (Gibco; #15240062), at 37°C in 5% CO₂. Cells were passaged every 2–3 days and used below passage 20.

### Generation of ISG15 Knockout Cell Lines

ISG15 knockout (KO) cell lines were generated by CRISPR/Cas9-mediated genome editing targeting exon 2 of the human ISG15 gene. Single-cell clones were isolated and validated functionally by immunoblotting for ISG15 expression following IFN-α or IFN-β stimulation, as indicated below.

#### Huh7 ISG15 KO cells

Huh7 ISG15 KO cell lines were generated using a transient plasmid-based Cas9 nickase strategy as previously described^35,36^. Guide RNAs targeting exon 2 of ISG15 were designed computationally and cloned into the Cas9 D10A nickase expression plasmid PX460 to minimize off-target editing^37^. Huh7 cells were transiently transfected with plasmids encoding paired sgRNAs and Cas9 nickase. Forty-eight hours after transfection, edited cells were enriched by puromycin selection and subsequently diluted to obtain single-cell clones. Cas9-expressing ISG15-intact clones generated in parallel were used as controls and are referred to throughout as wild-type (WT; Cas9 control) cells. Functional ISG15-knockout was confirmed by immunoblotting for ISG15 protein following IFN-α and IFN-β stimulation.

#### A549 ISG15 KO cells

A549 ISG15 KO cell lines were generated using a lentiviral CRISPR/Cas9 approach. The ISG15-targeting sgRNA sequence TTGAGGCCGTACTCCCCCAG was cloned into pLentiCRISPR v2 (GeneScript). Lentiviral particles were produced in Lenti-X 293 cells (Takara) by cotransfection of pLentiCRISPR v2-ISG15 sgRNA together with psPAX2 and pMD2.G using Lipofectamine 2000. Viral supernatants were collected and used to transduce A549 cells. Transduced cells were selected with puromycin (1 µg/mL), and individual clones were isolated. Functional knockout of ISG15 was confirmed by western blot analysis following IFN-α and IFN-β interferon stimulation.

#### Generation of HeLa-hACE2 and Subsequent ISG15 Knockout

HeLa-hACE2 cells were generated prior to ISG15 editing. HeLa cells were transduced with a VSV-G–pseudotyped lentivirus encoding pLVX-ACE2-BSD. Polyclonal cells were selected using blasticidin and subjected to limiting dilution cloning. Clones were screened for moderate ACE2 expression and efficient SARS-CoV-2 entry and replication; one clone was expanded for downstream experiments. ISG15 knockout in HeLa-hACE2 cells was subsequently introduced using ribonucleoprotein (RNP)-based CRISPR/Cas9 editing. Recombinant SpCas9-3xNLS was complexed with synthetic guide RNAs targeting exon 2 of ISG15 (sgRNA1: GGACCTGACGGTGAAGATGC; sgRNA2: CATGTCGGTGTCAGAGCTGA). RNP complexes were delivered by nucleofection (Lonza 4D-Nucleofector, pulse CM-137). Single-cell clones were isolated and validated by immunoblotting following IFN-β stimulation.

### TMT-Based Quantitative Proteomics by Mass Spectrometry

For quantitative proteomic analysis, Huh7 Cas9 control (ISG15-intact) and ISG15 knockout (KO) cells were seeded in 10-cm culture dishes and treated 24 h later with recombinant human IFN-β (1,000 U/mL) for 24 h or left untreated, as indicated in Figure 1A. Cells were washed with ice-cold PBS and pelleted by centrifuging in ice-cold PBS. Cell pellets were lysed in 100 µL of 8M urea/ 0.1% ProteaseMAX (Promega) in 0.1M Tris-HCl (pH 8.5) supplemented with protease and phosphatase inhibitors with ultrasonication. Lysates were cleared by centrifugation at 14,000 × *g* for 15 min at 4°C, and protein concentrations were determined using Micro BCA protein assay kit (Thermo Fisher Scientific), according to the manufacturer’s instructions. Equal amounts of protein (25 µg) from each biological replicate were subjected to reduction with dithiothreitol (DTT), alkylation with iodoacetamide, and digested with sequencing-grade trypsin (1:25 enzyme-to-protein ratio). Buffer was diluted to 1 M urea before in solution trypsin digestion. Peptides were sequentially cleaned on C18-Hypersep plate, dried by vacuum centrifugation, and resuspended for isobaric labeling.

Peptides were labeled using TMTpro™ 16-plex reagents (Thermo Fisher Scientific) according to the manufacturer’s protocol. Labeling efficiency (>95%) was confirmed prior to pooling. Labeled samples were combined in equal ratios and fractionated by high-pH reversed-phase chromatography using an offline C18 column. Fractions were concatenated into 12 pooled fractions to reduce sample complexity prior to LC-MS/MS analysis.

Each fraction was analyzed on an Ultimate 3000 UHPLC system (Thermo Fisher Scientific) coupled to an Orbitrap Q-Exactive HF mass spectrometer (Thermo Fisher Scientific). Peptides were separated on a reversed-phase analytical column using a 90-min linear gradient of increasing acetonitrile concentration in 0.1% formic acid. Data were acquired in data-dependent acquisition (DDA) mode. Full MS1 scans were collected at 120,000 resolution over an *m/z* range of 375–1,500 with a maximum injection time of 80 ms. The top 18 most intense precursor ions were selected for higher-energy collisional dissociation (HCD) fragmentation using a normalized collision energy of 34%, isolation width of 1.4 Th, dynamic exclusion of 45 s, and MS2 spectra acquired at 60,000 resolution with a maximum injection time of 54 ms.

Raw data were processed using Proteome Discoverer v2.4 (Thermo Fisher Scientific) with Sequest HT as the search engine. Spectra were searched against the UniProt SwissProt human database. Trypsin was specified as the protease with up to two missed cleavages allowed. Carbamidomethylation of cysteine was set as a fixed modification, while oxidation of methionine, deamidation (N/Q), and TMTpro modification of lysine residues and peptide N-termini were set as variable modifications. Peptide-spectrum matches were filtered using a 1% false discovery rate (FDR) at both peptide and protein levels. Relative protein abundances were calculated from TMT reporter ion intensities and normalized across channels prior to downstream statistical analysis.

### Immunoblotting

Cells were washed twice with ice-cold PBS and lysed on ice in NP-40 lysis buffer (50 mM Tris-HCl pH 7.5, 150 mM NaCl, 1% NP-40/Igepal CA-630, 1 mM EDTA) supplemented with protease inhibitor cocktail (cOmplete, Roche), phosphatase inhibitors (PhosSTOP, Roche), and 1 mM DTT where indicated. Lysates were incubated on ice for 30–50 min with intermittent vortexing and clarified by centrifugation at 14,000 × g for 15 min at 4°C. Protein concentrations were determined using the DC Protein Assay (Bio-Rad) according to the manufacturer’s instructions, using bovine serum albumin standards.

Equal amounts of total protein were mixed with 1× LDS sample buffer and 1× reducing agent, heated at 95-99°C for 10-15 min, and resolved (25–37.5 µg per lane) on 4–12% Bis-Tris NuPAGE gels (Thermo Fisher Scientific) in MOPS running buffer at 100 V for 30 min followed by 130 V for 90 min. Proteins were transferred to PVDF membranes using wet transfer in Tris–glycine buffer containing 20% methanol at 280 mA for 90 min at 4°C.

Membranes were blocked for 1 h at room temperature in 5% (w/v) non-fat dry milk in TBS-T (0.1% Tween-20) or, for phosphoprotein detection, in 5% (w/v) BSA in TBS-T. Primary antibodies were diluted 1:1,000 in blocking buffer and incubated either overnight at 4°C or for 2 h at room temperature with gentle agitation. Membranes were washed extensively in TBS-T and incubated with HRP-conjugated secondary antibodies (1:5,000) for 1–1.5 h at room temperature. After washing, signals were detected using enhanced chemiluminescence substrate and imaged using a digital imaging system. Exposure times were selected to ensure signal linearity.

Band intensities were quantified using ImageJ software. Target protein signals were normalized to loading controls (β-actin or Hsp60, as indicated). No membrane stripping was performed; when multiple proteins were analyzed on the same membrane, targets were selected based on sufficiently distinct molecular weights.

### RNA Extraction and Quantitative Real-Time PCR (qPCR)

Cell pellets were lysed in TRI reagent and homogenized by repeated pipetting at room temperature until complete disruption was achieved. RNA was purified using the Direct-zol RNA MiniPrep kit (Zymo Research), including on-column DNase I digestion to remove residual genomic DNA. RNA was eluted in RNase-free water, and concentration and purity were assessed by spectrophotometry (NanoDrop). RNA integrity was verified by A260/280 and A260/230 ratios. Equal amounts of total RNA were used for reverse transcription.

Complementary DNA (cDNA) was synthesized using the ABI High-Capacity cDNA Reverse Transcription Kit (Thermo Fisher Scientific) following the manufacturer’s protocol. Reverse transcription reactions were performed using equal RNA input amounts across samples, and resulting cDNA was diluted 1:5 in nuclease-free water prior to qPCR analysis.

Quantitative real-time PCR was performed using ABI Power SYBR Green Master Mix (Thermo Fisher Scientific) in 10 µL reactions containing diluted cDNA and 5 pmol of forward and reverse primers per reaction under the following cycling conditions: 95°C for 10 min, followed by 40 cycles of 95°C for 15 s and 60°C for 1 min. Melt curve analysis was performed to confirm specificity of amplification and absence of primer-dimer formation.

Infection with SARS-CoV-2 under BSL3 and CCHFV strain IbAr10200 under BSL4 condition followed the above-mentioned methodology following lysing the infected cells directly on plate with TRIzol. For infection with CCHFV-ZsG (strain IbAr10200) performed under BSL4 condition, total RNA was extracted using either the Magmax system (Life Technologies) or TRIzol/Direct-zol RNA MiniPrep (Zymo Research). One-step qRT-PCR was conducted using the SuperScript III Platinum One-Step qRT-PCR Kit (Life Technologies) and Taqman primer/probe sets (Thermo Fisher Scientific). Amplification of the CCHFV S segment followed previously described protocols^16^. Host cell transcripts (PPIA) were used for normalization, with all reactions performed on a Bio-Rad Thermal Cycler.

Relative gene expression was calculated using the comparative Ct (ΔΔCt) method. Ct values of target genes were normalized to the housekeeping gene HPRT, which was confirmed to be stable across experimental conditions. Fold changes were calculated as 2^−ΔΔCt relative to control samples.

### IFN-β Time-Course Stimulation

For kinetic analyses of IFN signaling and downstream ISG induction, Huh7 ISG15-intact (WT) and ISG15 knockout (KO) cells were seeded in 6-well plates at a density of 4 × 10^5^ cells per well in 2 mL high-glucose DMEM and allowed to adhere for 24 h. Cells were then stimulated with recombinant human IFN-β (1,000 U/mL). Parallel untreated control wells received fresh medium without IFN-β.

Cells were harvested at 3, 6, 12, and 24 h following IFN-β addition. For each experiment, an untreated control (0 h) was included and harvested at the final time point to ensure identical culture duration across conditions. At each time point, cells were washed once with ice-cold PBS and collected by scraping in ice-cold PBS. Cell pellets were obtained by centrifugation (5,000 × g, 5 min, 4°C) and processed for downstream analyses.

For immunoblotting, pellets were lysed in NP-40 lysis buffer supplemented with protease and phosphatase inhibitors as described above. For RNA analysis, parallel aliquots were processed for TRIzol-based RNA extraction and subsequent qPCR.

### Plasmids and ISG15 Overexpression

ISG15 overexpression experiments were performed in Huh7 Cas9-control (ISG15-intact) and ISG15 knockout (KO) cells using plasmids encoding HA-tagged human ISG15 with either an intact C-terminal di-glycine motif (ISG15-GG; conjugation-competent) or a mutated di-alanine motif (ISG15-AA; conjugation-deficient) in pcDNA3.1 vector backbone. An HA-empty vector served as control. All constructs were verified by Sanger sequencing prior to use.

Cells were seeded in 6-well plates at a density of 4 × 10^5^ cells per well in 2 mL high-glucose DMEM and allowed to adhere for 24 h at 37°C with 5% CO₂. Cells were transfected with 2.5 µg plasmid DNA per well using Lipofectamine 3000 and P3000 reagent (Thermo Fisher Scientific) according to the manufacturer’s protocol. After 24 h, medium was replaced and cells were stimulated with recombinant human IFN-β (1,000 U/mL) for an additional 24 h.

Cells were harvested by washing once with ice-cold PBS followed by lysis in NP-40 buffer (50 mM Tris-HCl pH 7.5, 150 mM NaCl, 1% NP-40, 1 mM EDTA) supplemented with protease and phosphatase inhibitors. Protein expression of ISG15 constructs and downstream targets was analyzed by immunoblotting as described above.

### Cycloheximide chase assay

Huh7 wild-type (WT) and ISG15 knockout (KO) cells were seeded in 6-well culture plates at a density of 0.4 × 10^6 cells per well in 2 mL of high-glucose DMEM. After 24 h of incubation at 37 °C in a humidified atmosphere containing 5% CO₂, cells were treated with 1,000 U/mL interferon-β (IFN-β) for 12 h.

Following IFN-β treatment, cells were exposed to cycloheximide (CHX) at a final concentration of 50 µg/mL (0.1% DMSO) or co-treated with CHX (50 µg/mL) and MG-132 (5 µM). To avoid replacing the IFN-β–containing medium, cycloheximide (100 mg/mL stock in DMSO) and MG-132 (10 mM stock in DMSO) were diluted in high-glucose DMEM to 2,050 µg/mL CHX and 205 µM MG-132, and 50 µL of these dilutions was added dropwise to wells containing 2 mL medium, resulting in the desired final concentrations. Control cells received an equivalent volume of DMSO to reach a final concentration of 0.1%.

CHX-treated cells were harvested at 0, 3, 6, and 12 h after CHX addition. Cells co-treated with CHX and MG-132, as well as DMSO-treated controls, were harvested after 12 h of treatment. At each harvest, cells were washed once with ice-cold PBS and collected on ice by scraping in 1 mL of ice-cold PBS. Cell suspensions were transferred to microcentrifuge tubes, pelleted at 5,000 × g for 5 min at 4 °C, and supernatants were removed. Pellets were stored at −80 °C until all conditions were collected. Cells were lysed in NP40 lysis buffer and protein levels were analyzed by Western blotting as described above.

### Immunoprecipitation

Huh7 wild-type (WT) and ISG15 knockout (KO) cells were seeded in 100 mm culture dishes at a density of 4 × 10^6 cells per dish in 8 mL DMEM. After 24 h of incubation at 37 °C in a humidified atmosphere containing 5% CO₂, cells were treated with 1,000 U/mL interferon-β (IFN-β) for 12 h.

Following IFN-β treatment, cells were exposed to MG-132 at a final concentration of 5 µM (0.05% DMSO). To avoid replacing the IFN-β containing medium, a 10 mM DMSO stock of MG-132 was diluted in DMEM to 205 µM, and 200 µL of this dilution was added dropwise to each culture dish containing 8 mL of medium, resulting in the desired final concentrations of MG-132 and DMSO. Control cells received an equivalent volume of DMSO to reach a final concentration of 0.05%.

After 12 h of MG-132 treatment (or 24 h of IFN-β treatment for control samples), cells were washed once with ice-cold PBS and lysed in 100 µL NP-40 lysis buffer (50 mM Tris-HCl pH 7.6, 150 mM NaCl, 1 mM EDTA, 1 mM DTT, 1% Igepal) supplemented with 1% sodium dodecyl sulfate (SDS). Prior to immunoprecipitation, 900 µL NP-40 lysis buffer was added to adjust the SDS concentration to 0.1%. The lysates were incubated with USP18 antibody for 2 h at 4 °C, followed by incubation with gamma-bind Sepharose beads (GE healthcare) for 2 h at 4 °C with rotation. The beads were washed with lysis buffer containing 0.1% SDS. Bound proteins were eluted by boiling in 2× SDS-PAGE loading buffer. Equal amounts of protein were subsequently analyzed by Western blotting as described above.

### RSAD2 (Viperin) Knockdown

For mechanistic interrogation of viperin function, RSAD2 was transiently silenced in Huh7, A549, and HeLa-hACE2 wild-type (WT) and ISG15 knockout (KO) cells using siRNA-mediated knockdown. Cells were transfected with a total of 54 pmol (6-well) or equimolar amounts scaled to dish surface area of siRNA targeting RSAD2, consisting of three independent siRNAs directed against distinct exons to minimize off-target effects. Control cells received an equal amount of non-targeting (scrambled) siRNA. Transfections were performed using Lipofectamine RNAiMAX (Thermo Fisher Scientific) according to the manufacturer’s protocol in antibiotic-free medium.

Six hours post-transfection, medium was replaced with complete DMEM, and cells were incubated for an additional 18 h before subsequent experimental manipulations. Where indicated, cells were treated with recombinant human IFN-β (1,000 U/mL) for 24 h prior to analysis. For viral infection experiments, siRNA-transfected cells were infected at the indicated multiplicity of infection following IFN-β pre-treatment when applicable. For metabolite quantification experiments, cells were harvested 24 h after IFN-β treatment by rapid washing in ice-cold PBS and extraction in pre-chilled 80% methanol. Knockdown efficiency was confirmed at both mRNA and protein levels by quantitative RT-PCR and immunoblotting prior to downstream analyses.

### Metabolite Extraction

Cells from 60 mm petri dishes were washed once with ice-cold PBS and directly scraped into 500 µL of ice-cold 80% methanol. Lysis was completed by probe sonication (10 s 20% amplitude, Qsonica Sonicator Q500) of dry-ice cold cell suspension. Following sonication, 200 µL of chloroform was added, and samples were vortexed for 5 s. Subsequently, 200 µL of Milli-Q water was added, followed by vortexing for 10 s and vigorous inversion for 30 s. Samples were centrifuged at 23,000 × g for 3 min at 0 °C, resulting in three distinct phases: an upper methanol-water phase containing polar metabolites, a lower chloroform–methanol phase, and a thin protein pellet at the interface. The upper phase was carefully collected for metabolite analyses. Methanol was removed by centrifugal vacuum evaporation for approximately 2 h. Protein fraction was recovered by mixing the chloroform phase and the interphase with 1 ml MeOH and centrifugation. The pelleted proteins were solubilized in 2% SDS and 0.5 % β-mercaptoethanol (in 60 mM Tris-Cl pH 6.8, 10% glycerol and 1 mM EDTA). Cellular protein content was quantified for normalization with a modified Bradford reagent, containing 2.5 mg/ml α-cyclodextrin to chelate excess SDS^38^, and albumin as a reference protein.

### Enzymatic Ribonucleotide Quantification

Cellular ribonucleoside triphosphate (rNTP) levels were quantified using an enzymatic microplate assay based on fluorogenic in vitro transcription^19^. Metabolite fractions were diluted to 1,200 µL per mg of cellular protein and briefly heat-denatured (3 min, 99 °C). Assay reactions (6–10 µL) were prepared in 384-well qPCR plates on ice and mixed with a 2X reaction mix, yielding final concentrations of 20 mM Tris-acetate (pH 7.9), 10 mM magnesium acetate, 50 mM potassium acetate, 0.1 mg/mL BSA, 2 mM spermidine, 1 mM of each non-limiting rNTP, 1 µM of the rNTP to be quantified, 20 nM DNA template encoding a dimeric stabilized Broccoli aptamer, 5 mM DTT, 20 µM BI (Broccoli aptamer ligand; BI), 0.5 kU/mL RiboLock RNase inhibitor, 0.25 U/mL pyrophosphatase, and 1.88 kU/mL T7 RNA polymerase (for ATP, UTP, and CTP quantification) or 0.5 kU/mL SP6 RNA polymerase (for GTP quantification).

Plates were sealed, briefly centrifuged, prewarmed on an aluminum block (37 °C, ∼30 s), and placed in a plate reader. Fluorescence was measured at 470 nm excitation and 510 nm emission at 37 °C until the reactions reached a plateau (2–3 h). Baseline-subtracted end-point fluorescence values were used to generate standard curves and determine sample rNTP concentrations, which were normalized to protein content of the initial sample.

### T7 RNA Polymerase Interference Test

To assess potential ddhCTP-mediated interference with the T7 RNA polymerase-based CTP assay, one 28 µL aliquot of the metabolite extract was mixed with 2 µL of water and another aliquot with 2 µL of 50 µM CTP (final added concentration 3.33 µM). CTP concentrations were measured using the enzymatic microplate assay described above, both with and without the added CTP, to determine reaction efficiencies. The Standard Addition Method was used to quantify interference in CTP quantifications.

### Nucleotide Quantification by HPLC

Intracellular nucleotides were quantified by isocratic reverse-phase HPLC with tetrabutylammonium as an ion-pairing agent as previously described^39^. In contrast to the original protocol, the UV detector was set to a wavelength of 280 nm.

### Virus infection

All experiments involving infectious Crimean-Congo hemorrhagic fever virus (CCHFV) were conducted under BSL4 biosafety containment conditions in accordance with institutional biosafety approvals. The CCHFV strain IbAr10200 (originally isolated from Hyalomma excavatum ticks in Nigeria, 1966) and recombinant CCHFV strain IbAr10200 expressing ZsGreen (CCHFV/ZsG) were generated and titrated as previously described^31,40^. For infections with the CCHFV isolate (IbAr10200), Huh7 wild-type (WT) and ISG15 knockout (KO) cells were seeded 24 h and transfected with Flag-Empty-CMV10 vector (for another study) prior to infection at multiplicity of infection (MOI 0.1), and virus adsorption was allowed to proceed for 1 h at 37°C. After adsorption, inoculum was removed, cells were washed once with PBS, and fresh complete medium (with or without IFN-β as specified) was added. Cells were lysed with TRIzol for downstream qPCR.

For fluorescence-based infection assays, A549 WT and ISG15 KO cells were infected with recombinant CCHFV/ZsG. ZsG fluorescence was monitored at 24-72 h post-infection (hpi) using a fluorescence plate reader or imaged using fluorescence microscopy. Fluorescence intensity was normalized to mock-treated infected controls to quantify relative viral replication.

CCHFV/ZsG infectious virus in supernatants was quantified by median tissue culture infective dose (TCID₅₀) assay using BSR-T7/5 cells. Serial ten-fold dilutions of supernatants were applied to BSR-T7/5 monolayers, and infection was detected either by monitoring ZsG fluorescence. TCID₅₀ values were calculated according to the Reed-Muench method.

For pharmacologic perturbation of the ISG15-USP18 axis, A549 WT and ISG15 KO cells were treated with ZHAWOC8655 at indicated concentrations. For infection experiments, cells were infected with CCHFV/ZsG, and ZHAWOC8655 was added post-infection at the indicated concentrations. DMSO concentration was kept constant across all conditions (≤0.025%). Viral replication was quantified by ZsG fluorescence and confirmed by TCID50 where indicated.

To assess compound cytotoxicity, parallel mock-infected cells were treated with identical concentrations of ZHAWOC8655 and cell viability was measured after 24 h using the CellTiter-Glo (Promega) luminescent ATP assay according to the manufacturer’s instructions. Luminescence values were normalized to DMSO-treated controls to determine relative viability.

All infections with SARS-CoV-2 were performed under BSL3 conditions. The SARS-CoV-2 isolate SWE/01/2020 (GenBank: MT093571) was originally recovered from a nasopharyngeal sample in Sweden and sequence-verified. Virus stocks were propagated in Vero E6 cells and titrated by TCID₅₀ assay on Vero E6 cells. HeLa-hACE2 wild-type and ISG15 knockout cells were seeded in 6-well plates and infected 24 h later with SARS-CoV-2 at a multiplicity of infection (MOI) of 0.1; viral RNA levels were quantified by qRT-PCR at the indicated post-infection time points.

For experiments assessing the role of viperin, RSAD2 knockdown was performed in A549 cells for CCHFV/ZsG, and HeLa-hACE2 cells for SARS-CoV-2 using siRNA transfection 24 or 48 h prior to infection. Cells were transfected with a pool of three siRNAs targeting independent RSAD2 exons (equal molar ratio) using Lipofectamine RNAiMAX according to the manufacturer’s instructions. Control cells received scrambled siRNA. Cells were subsequently infected with CCHFV/ZsG or SARS-CoV-2 as described above. Knockdown efficiency was validated by qPCR in parallel samples.

### CCHFV Minigenome assay

Huh7 wild-type (WT) and ISG15 knockout (KO) cells were seeded at a density of 0.03 × 10^6 cells per well in a 48-well plate. After 24 h, cells were transfected with 60 ng of pC-V5-L (or control vector), 20 ng of pC-N (or control vector in initial system testing), and 20 ng of pGL4.53 firefly control plasmids using 0.2 µL Lipofectamine 3000 reagent and 0.2 µL P3000 reagent (Thermo Fisher Scientific) per well, according to the manufacturer’s instructions. The condition in which pC-V5-L was transfected and pC-N was replaced with a control vector was only included in the initial testing of the minigenome system, but was not included in experiments incorporating IFN-β treatment or RSAD2 knockdown.

Twenty-four hours after plasmid transfection, the medium was replaced with fresh medium, and cells were transfected with 100 ng of T7-minigenome (T7-MG) in vitro transcripts using 0.2 µL Lipofectamine MessengerMAX Transfection Reagent (Thermo Fisher Scientific) per well. Four hours after T7-MG RNA transfection, the medium was replaced with fresh medium. Sixteen hours after T7-MG RNA transfection, cells were washed once with ice-cold PBS and lysed in 80 µL of 1X Passive Lysis Buffer (Promega) for 45 min at room temperature on an orbital shaker. Lysates were mixed by pipetting, and 60 µL was transferred to wells of a white, flat-bottom 96-well Opti-Plate (Corning). Firefly and NanoLuc luminescence were measured using a Tecan plate reader with the Nano-Glo Dual-Luciferase Reporter Assay System (Promega) according to the manufacturer’s instructions for Passive Lysis Buffer.

For IFN-β–treated experiments, 4 h after plasmid transfection the medium was replaced with fresh medium. Sixteen hours after plasmid transfection, the medium was removed and cells were treated with 1,000 U/mL IFN-β. Eight hours after IFN-β treatment (24 h after plasmid transfection), T7-MG RNA transfection proceeded as described above.

For experiments including RSAD2 knockdown, 24 h after seeding, Huh7 WT and ISG15 KO cells were transfected with 2.5 pmol total siRNA comprising equal amounts of three different siRNAs targeting three different exons of RSAD2 complexed with 0.75 µL Lipofectamine RNAiMAX Transfection Reagent (Thermo Fisher Scientific) per well, according to the manufacturer’s instructions. Control cells were transfected with 2.5 pmol of scrambled siRNA per well. Six hours after siRNA transfection, the medium was replaced with fresh medium, and experiments continued with plasmid transfection, followed by IFN-β treatment and T7-MG RNA transfection as described above.

### ZHAWOC8655 Treatment and Functional Assays

For pharmacologic perturbation of the ISG15-USP18 axis, A549 wild-type (WT) and ISG15 knockout (KO) cells were seeded in 6-well plates at 0.4 × 10⁶ cells per well in 2 mL high-glucose DMEM and allowed to adhere for 24 h at 37°C with 5% CO₂. Where indicated, cells were stimulated with recombinant human IFN-β (1,000 U/mL) concurrently with ZHAWOC8655 treatment. ZHAWOC8655 was added at final concentrations of 6.25 µM or 12.5 µM; 25 µM was excluded from infection experiments due to toxicity observed under virus infection conditions. All treatments contained 0.025% DMSO as vehicle control.

For immunoblot analysis, cells were treated for 24 h, washed with ice-cold PBS, and lysed in NP-40 lysis buffer supplemented with protease and phosphatase inhibitors. Lysates were clarified by centrifugation and processed for SDS-PAGE and immunoblotting as described above.

For CCHFV infection experiments, A549 WT cells were infected under standard infection conditions as described in the virus infection section. ZHAWOC8655 was added after viral adsorption, and infection proceeded for 24 h in the continued presence of compound. Viral replication was quantified by the indicated readouts (ZsGreen fluorescence, viral RNA quantification, or TCID₅₀), depending on the experiment. Only non-toxic concentrations were included in quantitative analyses.

Cell viability under compound treatment was assessed using the CellTiter-Glo Luminescent Cell Viability Assay (Promega). A549 WT cells were seeded in 96-well plates at 1 × 10⁴ cells per well in 100 µL DMEM. After 24 h, cells were treated with IFN-β (1,000 U/mL) in combination with ZHAWOC8655 (0-25 µM) or vehicle control (0.025% DMSO). After 24 h, luminescence was measured according to the manufacturer’s protocol.

## Supplementary Figures

**Figure S1. ISG15 deficiency reshapes the interferon transcriptional landscape toward amplified antiviral effector and immune signaling programs (Related to Figure 1 and Figure 2). (A)** Unsupervised hierarchical clustering heatmap of differentially expressed genes across WT and ISG15KO cells ± IFN-β stimulation. Gene expression values are Z-score normalized. Samples cluster primarily by IFN treatment and genotype, with ISG15KO + IFN samples forming a distinct transcriptional module characterized by exaggerated induction of interferon-stimulated genes (ISGs). The comparison bar indicates genes differentially expressed in KO+ vs KO− and WT+ vs WT− conditions. Color scale represents relative expression (yellow, high; blue, low). **(B)** Volcano plot of differentially expressed genes in WT cells following IFN-β stimulation (WT+ vs WT−). The x axis indicates log2 fold change; the y axis shows –log10 adjusted p value. Bubble size reflects absolute log2 fold change. Canonical ISGs including RSAD2, OAS family members, MX1, IFITM3, and BST2 are significantly upregulated. The magnitude of induction in WT cells is robust but constrained relative to KO cells (see panel C). **(C)** Volcano plot of differentially expressed genes in ISG15KO cells following IFN-β stimulation (KO+ vs KO−). Compared to WT, KO cells exhibit enhanced amplitude and breadth of ISG induction, with increased fold changes and statistical significance across antiviral effectors, including OAS2, SP100, IFI6, DDX60, ISG15, IFITM3, BST2, TAP2, and HLA genes. The expanded transcriptional dynamic range supports loss of negative feedback regulation. **(D)** Gene set enrichment analysis (GSEA) identifying pathways uniquely enriched in WT cells following IFN stimulation. Enriched pathways include E2F targets, mTORC1 signaling, heme metabolism, TNFα signaling via NF-κB, KRAS signaling, inflammatory response, complement, coagulation, adipogenesis, and G2/M checkpoint. Dot size represents pathway overlap; color intensity indicates –log10 adjusted p value. These pathways suggest a balanced IFN response integrating immune signaling with cell-cycle and metabolic control. **(E)** GSEA identifying pathways uniquely enriched in ISG15KO cells following IFN stimulation. KO cells show strong enrichment of interferon alpha and gamma responses, IL-6/JAK/STAT3 signaling, TNFα signaling, p53 pathway, apoptosis, hypoxia, allograft rejection, TGF-β signaling, epithelial–mesenchymal transition, cholesterol homeostasis, peroxisome, and PI3K/AKT/mTOR signaling. The magnitude and diversity of enriched immune and stress pathways indicate loss of transcriptional restraint and amplified inflammatory signaling. **(F)** Global distribution of differentially expressed genes (KO+ vs KO−) highlighting the expanded dynamic range of ISG induction in ISG15-deficient patient derived fibroblast cells± IFN-α stimulation from reanalysis and replotting of previously published transcriptomic data to illustrate conserved induction patterns of select ISGs (data adapted from Akalu et al.^12^ Sci Transl Med. 2025). This external dataset is included for reference and was not generated in the current study. Bubble size corresponds to absolute log2 fold change; y axis indicates –log10 adjusted p value. The genes also found in the proteomics data is highlighted in bright color and the genes not observed in the proteomics are in faded shade. Notably, RSAD2, OAS2, BST2, CMPK2, and MX1 rank among the most strongly induced transcripts in the current dataset.

**Figure S2. ISG15 deficiency prolongs STAT1/STAT2 activation and enhances downstream ISG transcription and protein accumulation. (Related to Figure 2). (A–D)** Quantitative analysis of proximal JAK–STAT signaling following IFN-β (1000 U/mL) stimulation (0–24 h) in WT and ISG15KO cells. Immunoblot signals were normalized to β-actin and expressed relative to WT at 24 h. **(A)** Phospho-STAT2 (p-STAT2). **(B)** Total STAT2. **(C)** Phospho-STAT1 (p-STAT1). **(D)** Total STAT1. ISG15KO cells exhibit significantly enhanced and prolonged STAT1 and STAT2 phosphorylation compared to WT, whereas total STAT1 and STAT2 levels show only modest differences, indicating dysregulated signaling amplitude rather than altered STAT abundance. **(E–H)** Quantification of downstream ISG protein accumulation following IFN-β stimulation (0–24 h). Protein levels were normalized to β-actin and expressed relative to WT at 24 h. **(E)** IRF9. **(F)** MX1. **(G)** IFIT1. **(H)** Viperin (RSAD2). ISG15KO cells display amplified late-phase accumulation of antiviral effector proteins, with RSAD2 showing the most pronounced divergence from WT. When C_T_ values were above 35 (undetectable) at the 0h time-point, the same delta C_T_ value was set for WT and KO at the 0h time-point only. Data represent mean ± SEM from independent biological replicates (n ≥ 3). Statistical analysis was performed using two-way ANOVA with multiple comparisons; ns, not significant; *p < 0.05; **p < 0.01; ***p < 0.001; ****p < 0.0001. **(I–L)** Time-course analysis of ISG mRNA induction following IFN-β treatment (0–24 h). Transcript levels were normalized to HPRT and expressed as fold change relative to WT 0 h (CMPK2) or WT 24 h (MX1/IFIT1/RSAD2). **(I)** CMPK2 mRNA. **(J)** MX1 mRNA. **(K)** IFIT1 mRNA. **(L)** RSAD2 mRNA. ISG15KO cells demonstrate enhanced transcriptional output at later time points, particularly for RSAD2, consistent with sustained STAT activation translating into amplified gene expression. Data represent mean ± SEM from independent biological replicates (n ≥ 3). Statistical analysis was performed using two-way ANOVA with multiple comparisons; ns, not significant; *p < 0.05; **p < 0.01; ***p < 0.001; ****p < 0.0001. **(M–N)** Rescue of sustained STAT phosphorylation by ISG15 reconstitution. ISG15KO cells were transfected with empty vector (EV), HA-ISG15-GG (conjugation-competent), or HA-ISG15-AA (conjugation-defective) for 24 h followed by IFN-β stimulation (1000 U/mL, 24 h). **(M)** Quantification of p-STAT1 normalized to β-actin. **(N)** Quantification of p-STAT2 normalized to β-actin. Reintroduction of conjugation-competent ISG15 significantly attenuates prolonged STAT phosphorylation, whereas the conjugation-defective mutant fails to fully restore regulation. **(O)** Quantification of viperin protein levels following ISG15 reconstitution. Viperin abundance was normalized to HSP60 and expressed relative to KO EV + IFN. Conjugation-competent ISG15 significantly suppresses viperin accumulation, whereas ISG15-AA shows only partial rescue, supporting a conjugation-dependent role for ISG15 in restraining amplified antiviral effector output. Data represent mean ± SEM from independent biological replicates (n ≥ 3). Statistical analysis was performed using ordinary one-way ANOVA; ns, not significant; *p < 0.05; **p < 0.01; ***p < 0.001; ****p < 0.0001.

**Figure S3. Reduced CCHFV RNA accumulation and reporter virus signal in ISG15-deficient cells. (Related to Figure 4). (A)** Quantification of CCHFV S-segment RNA levels in Huh7 WT and ISG15KO cells following infection with CCHFV (strain IbAr10200). Viral RNA was measured by qRT-PCR, normalized to HPRT, and expressed as fold change relative to WT mock-infected cells. ISG15KO cells exhibit significantly reduced viral RNA accumulation compared to WT cells under identical infection conditions. Data are shown as mean ± SEM from independent biological replicates (n ≥ 3). Statistical analysis was performed using one-way ANOVA; ****p < 0.0001. **(B)** RSAD2 mRNA expression in WT and ISG15KO cells following CCHFV infection. Transcript levels were normalized to HPRT and expressed relative to WT mock-infected controls. ISG15KO cells display increased RSAD2 induction upon infection compared to WT, consistent with amplified IFN signaling and enhanced antiviral effector expression. **(C)** Replication kinetics of recombinant CCHFV/ZsGreen (ZsG) reporter virus (MOI 0.1) in A549 WT and ISG15KO cells. Viral replication was monitored by ZsG fluorescence at 0, 24, 48, and 72 h post-infection. ISG15KO cells demonstrate reduced reporter signal at later time points compared to WT cells, indicating attenuated viral replication. Data represent mean ± SEM from independent biological replicates (n ≥ 3). Statistical analysis was performed using two-way ANOVA with multiple comparisons; ns, not significant; ****p < 0.0001.

**Figure S4. ddhCTP accumulation in ISG15-deficient cells is viperin-dependent. (Related to Figure 3). (A)** Quantification of intracellular ribonucleotide triphosphate (rNTP) levels in WT and ISG15KO cells ± IFN-β stimulation. ATP, GTP, UTP, and CTP levels were measured by HPLC and normalized to untreated WT controls. ATP and GTP levels are largely unchanged across genotypes and treatment conditions, whereas UTP and particularly CTP levels are reduced in ISG15KO cells and remain unchanged following IFN stimulation. Data represent mean ± SEM from independent biological replicates (n ≥ 3). **(B)** Absolute quantification of CTP and ddhCTP levels (µM) by HPLC in WT and ISG15KO cells ± IFN-β stimulation. ISG15KO cells display reduced CTP levels accompanied by increased ddhCTP accumulation upon IFN treatment, supporting enhanced viperin-mediated catalytic conversion of CTP. **(C)** Immunoblot validation of viperin (RSAD2) expression following siRNA-mediated knockdown (si-RSAD2) in WT and ISG15KO cells ± IFN-β stimulation. Hsp60 serves as loading control. Efficient depletion of viperin confirms specificity of downstream nucleotide measurements. The analyzed proteins are derived from the protein pellet after chloroform/methanol metabolite extraction of samples that were used for T7 polymerase activity and ddhCTP/CTP quantification (Figure 3 E-F). Immunoblots are representative of independent experiments (n ≥ 3) **(D)** Quantification of intracellular CTP levels (pmol/mg protein) using an enzymatic microplate assay based on fluorogenic *in vitro* transcription in WT and ISG15KO cells treated with si-scr or si-RSAD2 ± IFN-β. IFN-induced CTP depletion in ISG15KO cells is significantly restored upon RSAD2 knockdown, aligning with the reduced T7 polymerase activity. Data represent mean ± SEM from independent biological replicates (n ≥ 3). Statistical analysis was performed using ordinary one-way ANOVA; **p < 0.01; ****p < 0.0001. **(E)** Quantification of CTP and ddhCTP levels (µM) by HPLC following RSAD2 knockdown. Elevated ddhCTP accumulation observed in ISG15KO + IFN cells is significantly reduced upon viperin silencing, accompanied by restoration of intracellular CTP levels. Data represent mean ± SEM from independent biological replicates (n ≥ 3).

**Figure S5. RSAD2 knockdown restores CCHFV replication in ISG15-deficient cells and validation of the CCHFV minigenome system. (Related to Figure 4). (A)** Quantification of infectious CCHFV titers (TCID50/mL) in WT and ISG15KO cells following siRNA (si-scr or si-RSAD2) treatment. ISG15KO cells display reduced viral titers compared to WT under si-scr conditions, and RSAD2 knockdown significantly restores infectious virus production, confirming viperin-dependent restriction. Data represent mean ± SEM from technical replicates (n ≥ 3). Statistical analysis was performed using ordinary one-way ANOVA; **p < 0.01; ***p < 0.001. **(B)** Validation of in vitro-transcribed CCHFV minigenome (MG) RNA used in polymerase assays. Agarose gel electrophoresis demonstrates the integrity of T7-generated MG transcripts and removal of residual plasmid DNA following Turbo DNase treatment. **(C)** Specificity controls for the CCHFV minigenome assay. NanoLuc/Firefly luminescence ratios are shown under conditions lacking L polymerase (−Lpol) or N protein (−Nprot). NanoLuc/Firefly signal was normalized to (+Lpol) (+Nprot). Polymerase activity is observed only in the presence of L polymerase, confirming assay specificity and absence of background reporter activity. Data represent mean ± SEM from technical replicates (n ≥ 3).

**Figure S6. Pharmacologic disruption of the ISG15–USP18 axis enhances IFN signaling without overt cytotoxicity. (Related to Figure 4). (A)** Assessment of 24h cell viability following treatment with the USP18-ISG15 interaction inhibitor ZHAWOC8655 (0-25 µM) as indicated in presence of IFN-β (1000u/mL). CellTiter-Glo (CTG) luminescence signal is expressed as percentage relative to IFN-β treated but ZHAWOC8655 untreated controls (0 µM). ZHAWOC8655 induces a modest, dose-dependent reduction in viability, with >75–85% viability maintained at concentrations used for immunoblot experiments. Data represent mean ± SEM from technical replicates (n ≥ 3). **(B)** Immunoblot analysis of WT and ISG15KO cells treated with IFN-β (1000 U/mL) targeting ISGs in the presence of increasing concentrations of ZHAWOC8655 (0–25 µM). USP18, MX1, IFIT1, global ISGylation, and ISG15 levels are shown; β-actin serves as loading control. ISG15KO cells serve as a control for ISG15-dependent effects. The ISGylation smear confirms intact conjugation machinery in WT cells and absence in KO cells. **(C-E)** Quantification of protein levels of **(C)** USP18, **(D)** MX-1 and **(E)** IFIT1 from panel **(B)**, normalized to β-actin and relative to the IFN-β treated and ZHAWOC8655 non-treated WT cells. Immunoblot image and quantification are representative of two independent biological replicates.

## Supplementary Tables

**Table S1:** Quantitative TMT-based proteomic analysis of IFNβ-treated WT and ISG15 knockout Huh7 cells.

## Acknowledgements

This work was supported by the Swedish Research Council (Vetenskapsrådet) through a Starting Grant awarded to S.G. (2021–03035). Additional funding was provided by the Center for Medical Innovation (CIMED-FoUI093304) and Karolinska Institutet Stiftelser och Fonder (grant 2022–02232) to S.G. This work was also supported in part by CDC Emerging Infectious Disease Research Core funds and the National Institute of Allergy and Infectious Diseases (NIAID) grant R01AI151006 to E.B.. S.R. was supported by the Karolinska Institutet KID funding program awarded to S.G. J.P. was supported by Sigrid Jusélius Foundation and Karolinska Institutet Stiftelser och Fonder. U.N. acknowledge support received from European Union’s HORIZON Research and Innovation Actions under grant agreement No. 101191315 (COMBAT). U.N. also acknowledge support received from the Swedish Research Council grants (2021-01756, 2021-00993, 2018-06156 and 2017-01330), Karolinska Institutet Consolidator Grant (2-117/2023). We thank Hannah Nor and Annika Berggren (Uppsala University) for technical assistance during their laboratory training.

## Disclaimer

The findings and conclusions in this report are those of the authors and do not necessarily represent the official position of the Centers for Disease Control and Prevention. Views and opinions expressed are those of the authors only and do not necessarily reflect those of the European Union. The European Union cannot be held responsible for them.

## Author Contributions

Conceptualization: S.G., J.P., E.B.

Methodology: N.L.K., S.R., F.E.M.S., J.P., E.B., S.G., A.V., P.N., A.H., E.S., M.L.

Investigation: N.L.K., S.R., F.E.M.S., R.M., S.F., J.N., V.M.M., E.S.

Formal Analysis: A.T.A., A.H., A.V., P.N., R.L.-J.

Resources: S.G., E.B., C.G.G.B., V.P., Y.T.B., A.M., U.N., R.L.-J.

Supervision: S.G., J.P., E.B., Y.T.B., U.N., A.V., A.M.

Writing - Original Draft: S.G., N.L.K., S.R., J.P., F.E.M.S, E.B.

Writing - Review & Editing: N.L.K., S.R., F.E.M.S., J.P., E.B., A.T.A., U.N., C.G.G.B., A.V., A.H.

Funding Acquisition: S.G., E.B., J.P., U.N.

